# Time-resolved operator archetypes characterize dynamical sensitivity during cell-state transitions

**DOI:** 10.64898/2026.08.21.745996

**Authors:** David Redd, Sam Green, Tommy W. Terooatea

## Abstract

During development, cells traverse gene expression states where their local dynamical sensitivity changes sharply, yet existing computational methods provide limited access to when and where this sensitivity peaks along a trajectory. Here we introduce scJDO (single-cell Jacobian Differential Operators), a framework that characterizes how local dynamical sensitivity evolves during cell fate transitions, together with an explicit account of what that representation can and cannot recover from snapshot data.

scJDO treats time-indexed Jacobians as explicit analytical objects, projecting the temporal sequence of operators into a shared subspace and decomposing it into recurrent operator archetypes with interpretable temporal activation profiles. Unlike methods that derive Jacobians from splicing-kinetic vector fields, scJDO learns a neural drift field directly from cell-state geometry via diffusion score matching, enabling Jacobian analysis on trajectory-resolved scRNA-seq datasets regardless of splicing-data availability. Applied to a dense time-course of induced pluripotent stem cell (iPSC) reprogramming, scJDO resolves a quantitative operator-level signature that distinguishes diverted from productive fate: the productive trajectory executes a sequential handoff from an early MEF-exit operator regime to a late pluripotency-associated regime, whereas the diverted trajectory maintains the early regime and instead activates a distinct stress-associated archetype.

We validate scJDO across four settings: synthetic benchmarks with analytically known ground truth, branching hematopoiesis, dense real time-course reprogramming, and a perturbational setting using Schrödinger bridges in K562 CRISPRi Perturb-seq. We compare against the two most widely used single-cell Jacobian methods on a dataset where all three are runnable, finding that scJDO shares significantly more gene-level and directional operator structure with Dynamo than expected by chance while providing operator-level analysis on datasets without splicing kinetics. We further characterize the boundary of the representation directly. At a fate-decision saddle, eight mathematically distinct readouts of the same learned drift field are consistent with a single explanation: a drift field fit to snapshot density reproduces density-dominant separation between committed branches rather than the low-variance transverse instability that defines the decision. Together, scJDO provides an operator-level view of single-cell dynamics and an explicit characterization of its own identifiability boundary.

## Introduction

Cellular identity emerges through dynamic regulatory programs that evolve over developmental trajectories. As cells traverse high-dimensional state spaces, transitions in identity arise through coupled activity of transcription factors, signaling pathways, and chromatin-associated mechanisms. Single-cell RNA sequencing (scRNA-seq) has transformed our ability to observe these processes, yielding high-resolution snapshots of cellular heterogeneity and developmental progression. However, a critical question remains: where along a developmental trajectory does the local dynamical sensitivity of the system peak? Identifying these transient windows is relevant to optimizing directed differentiation protocols, timing drug interventions, and understanding the molecular drivers of fate decisions.

Here we introduce scJDO (single-cell Jacobian Differential Operators), a framework designed to identify when and where local dynamical sensitivity changes along a differentiation trajectory. Applied to a dense time-course of iPSC reprogramming, scJDO reveals that productive and diverted fates — arising from the same starting cells — are distinguished by a quantitative operator signature: the productive trajectory executes a sequential operator handoff, while the diverted trajectory arrests at an early program and activates a stress-associated archetype.

To achieve this, scJDO introduces an operator-centric view of cell fate dynamics. Current computational approaches — including pseudotime inference [1–3], RNA velocity [4,5], and probabilistic fate mapping [6] — have provided critical insight into trajectory geometry and fate probabilities, but largely operate at the level of where cells are likely to go rather than how local dynamical sensitivity evolves. From a dynamical-systems perspective, local properties such as stability and perturbation sensitivity are governed by the Jacobian of the underlying drift field. While local Jacobian analysis is central in dynamical systems, existing single-cell implementations (Dynamo [7], SpliceJAC [8]) derive Jacobians from splicing-kinetic vector fields, which require metabolic labeling or explicit assumptions about RNA production and degradation. Furthermore, they treat each Jacobian as a per-state regulatory snapshot.

scJDO addresses these limitations through two advances. First, it treats the temporal sequence of Jacobians as a primary analytical object: it projects the entire sequence into a shared lower-dimensional operator space via semi-nonnegative matrix factorization (semi-NMF) [9] and identifies recurrent archetypes of instability within that space. By stacking operators across pseudotime and decomposing the resulting tensor, scJDO identifies reusable local dynamical regimes and estimates when they become active, sequentially hand off, or recede. Second, unlike methods that derive Jacobians from splicing kinetics, scJDO learns a neural drift field directly from cell-state geometry via diffusion score matching [10,11], enabling Jacobian analysis on trajectory-resolved scRNA-seq datasets regardless of splicing-data availability.

Importantly, we do not position scJDO as a competitor to fate-mapping or velocity methods, which answer different questions. We show instead that, where it computes a comparable object — a local Jacobian — scJDO shares significantly more gene-level and directional structure with Dynamo than expected by chance, supporting the interpretation that the learned operators reflect data-driven biology rather than artifacts of the neural drift field, while yielding a distinct operator-level ordering not recovered by either splicing-derived method. We also delineate, against analytic ground truth and across our biological datasets, the conditions under which the method’s outputs are and are not identifiable. As established by Weinreb et al. [12], a snap-shot fixes the density of cell states but not the velocity field that generated it; we argue this boundary is a property of the snapshot data modality shared by all such methods, not of scJDO specifically. We characterize one specific facet of this boundary — instability at a bifurcation saddle — directly, with a dedicated Results subsection that reports what fails across eight mathematically distinct readouts of the same drift field, identifies a density-dominance mechanism, and reports a later-window density-enrichment control that does not rescue saddle localisation.

We test this framework across four validation settings — synthetic benchmarks with known ground truth, branching hematopoiesis [1], dense real time-course reprogramming [13], and a perturbational setting using Schrödinger bridges [14] in K562 [15] — and characterize the identifiability boundary of the operator representation throughout. In this way, scJDO provides a framework for studying when cell states appear dynamically buffered, plastic, or sensitive to perturbation, and an explicit account of the resolution at which those distinctions are supported.

## Results

### scJDO infers time-resolved operator representations of local sensitivity

The scJDO framework transforms static single-cell measurements into an operator-centric representation of cellular dynamics through four stages (Fig. 1). scJDO’s operator inference is performed independently and does not require predefined basins.

**Figure 1.**
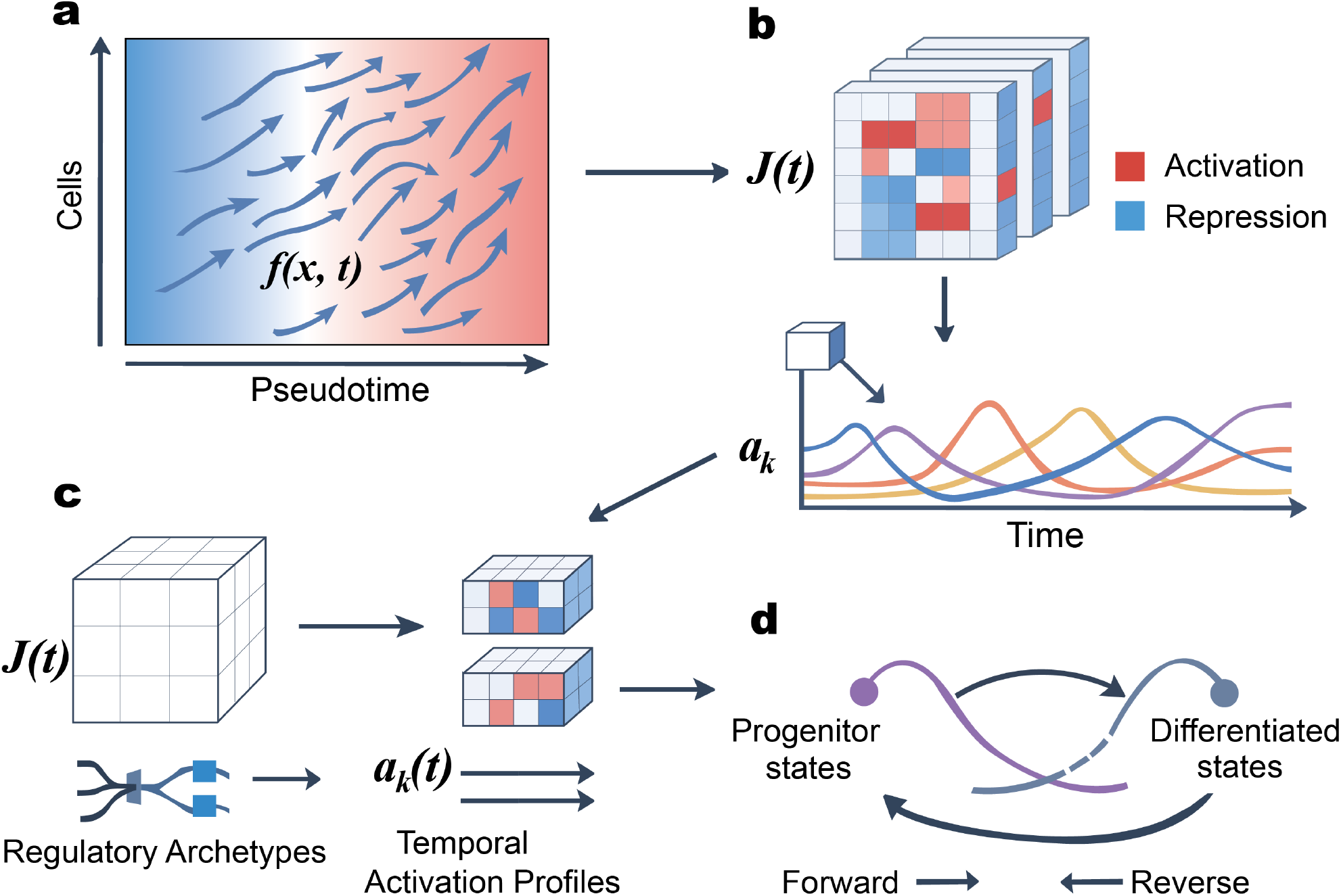
The scJDO framework. a, Drift-field estimation. A neural drift field f(x, τ) is learned from cell-state geometry in a latent space via diffusion score matching, with an ordering-consistent residual head and optional weak RNA-velocity guidance (Methods). Arrows indicate the inferred drift; the color gradient indicates progression along the supplied temporal ordering τ. b, Local operator inference. Local Jacobians J(τ) = ∂f/∂x are computed by automatic differentiation at each cell and aggregated across the ordering by Gaussian kernel weighting, giving a temporal Jacobian tensor of shape T × d × d. Matrix entries summarize local linearized sensitivity: positive (red) and negative (blue) couplings between latent directions, interpretable in gene space after projection through the FA loadings. c, Archetype decomposition. The tensor is unfolded along the time axis and factorized by semi-NMF into K operator archetypes A_k recurrent Jacobian regimes — with non-negative temporal activation profiles a_k(τ). Archetypes are recovered up to a column permutation of the factorisation, so component labels are arbitrary and only their temporal ordering is interpretable (Box 1). d, Trajectory synthesis. Activation profiles are read as a sequence of local dynamical regimes along the trajectory: which regimes are active, when they hand off, and where sensitivity peaks. For datasets without a natural pseudotime, the Schrödinger-bridge formulation computes forward and reverse operators between defined source and target populations (Fig. 6).

#### Stage 1: Drift-field estimation

We model cellular state evolution in a latent space as a stochastic differential equation (Fig. 1a):

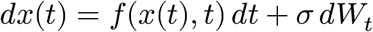

where *x*(*t*) ∈ ℝ^*d*^ is the latent cellular state, *f* (⋅, *t*) is a learned drift field, and *W*_*t*_ is a Wiener process. The drift *f* is parameterized by a hybrid architecture combining diffusion score matching [10,11] with residual deterministic components and optional RNA-velocity guidance, yielding a smooth vector field suitable for operator analysis (Fig. 1a; Methods). When used, RNA velocity acts as a weak directional prior on *f* and does not define macrostates, terminal basins, or archetypes.

#### Stage 2: Local operator inference (Fig. 1b)

We compute local Jacobian operators along inferred progression by automatic differentiation:

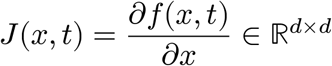

These operators provide a comparative summary of local linearized sensitivity — how small perturbations are predicted to be amplified or buffered around a point in state space (Fig. 1b). They arise from derivatives of the learned drift field, reflecting inferred local response properties rather than static covariance structure.

#### Stage 3: Temporal tensor construction (Fig. 1c)

We estimate a continuous operator trajectory by aggregating per-cell Jacobians using Gaussian kernel weighting along pseudotime, with bandwidth selected to balance instability-peak localization, bootstrap reproducibility, and a minimum effective sample size. The resulting smoothed operators are stacked into a temporal Jacobian tensor *J* ∈ ℝ^*T*×*d*×*d*^, where *T* indexes pseudotime grid points and *d* is the latent dimension. Temporal resolution is limited by local pseudotime density.

#### Stage 4: Archetype decomposition (Fig. 1d)

To identify recurrent operator regimes, we project the temporal sequence of Jacobians into a shared operator subspace. The tensor is unfolded along the time axis into a matrix 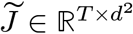 and decomposed via semi-NMF [9] (with SVD as a fallback), with decomposition rank selected as described in Methods; *K* = 5 was used for the biological branch analyses:

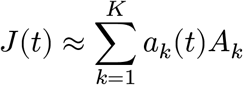

where *A*_*k*_ ∈ ℝ^*d*×*d*^ are operator archetypes — recurrent Jacobian regimes — and *a*_*k*_(*t*) are their time-varying activation profiles. This decomposition reveals both the repertoire of recurrent local dynamical regimes and the temporal patterns by which they sequentially hand off. Semi-NMF is ambiguous up to any invertible transformation *Q* such that *W Q* ≥ 0 (a cone ambiguity, not just permutation and scale). Despite this broader formal ambiguity, independently fitted solutions empirically converge to highly similar matched archetypes on synthetic ground truth (mean per-archetype cosine similarity on matched *H* pairs of 0.81 on sharp-step handoffs and 0.87 on sigmoid handoffs; see §Synthetic benchmarks and Fig. 2 legend), which bounds how much the cone ambiguity is exercised in practice; downstream comparisons therefore use explicit matching and interpret archetype identity operationally up to permutation rather than claiming uniqueness. Because component labels are arbitrary under this permutation, only the temporal ordering of archetypes — not their indices — carries interpretable content.

**Figure 2.**
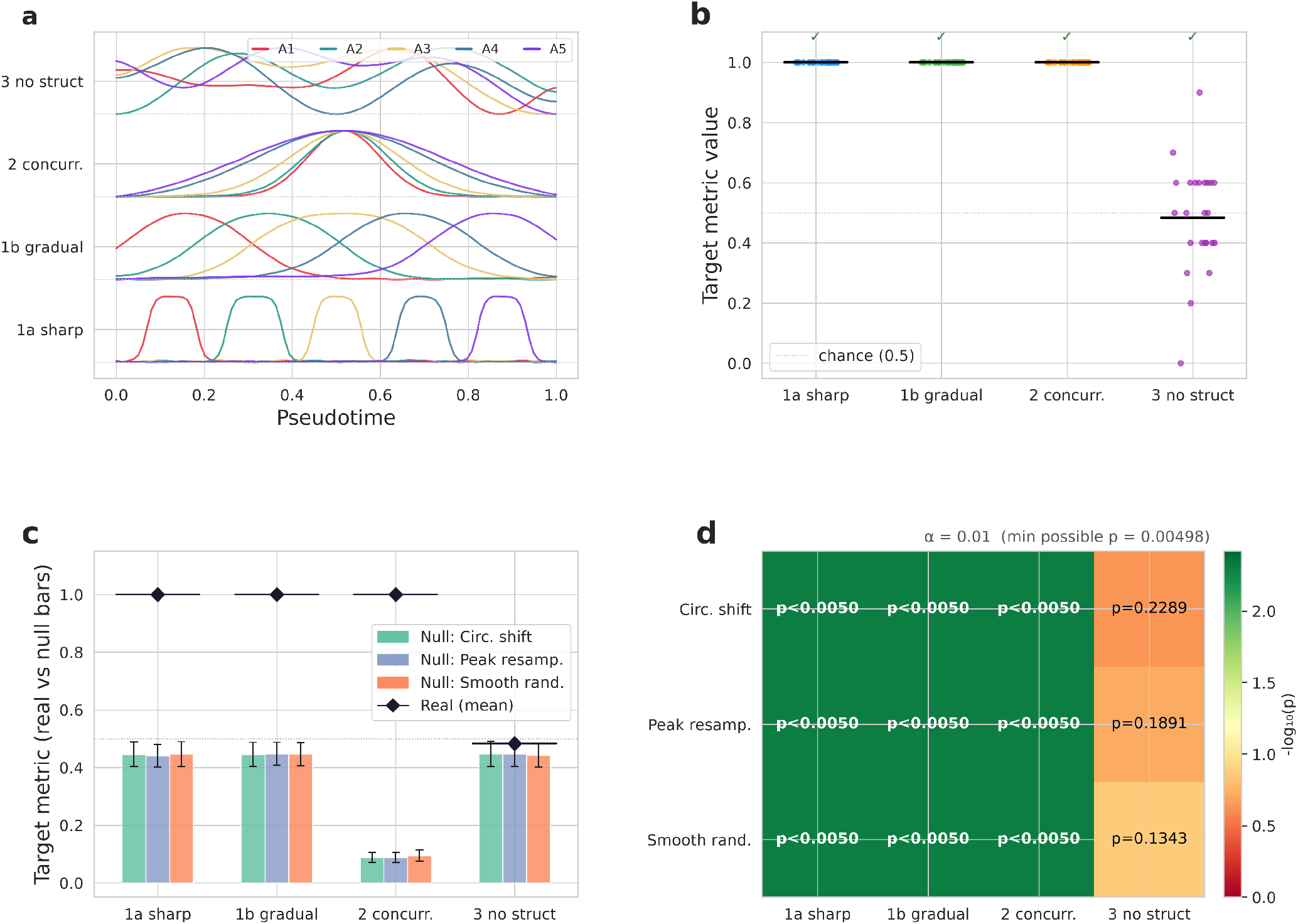
Peak-timing coordination metric — synthetic positive control. Four synthetic systems with five archetypes each: **1a** sharp gated pulses, **1b** sigmoid handoffs, **2** overlapping Gaussians co-timed at *τ* = 0.5, **3** unstructured. **(a)** Representative activation profiles per system. **(b)** Target metric across 24 seeds per system, approaching 1 on 1a/1b/2 (✓) and ≈ 0.5 (chance) on 3. **(c)** Real means (♦) versus three null-model means (bars, ±SD; circular-shift, peak-resampling, smooth-randomisation nulls) per system. **(d)** − log_10_ empirical *p*-value heatmap: all coordinated systems reach *p* < 0.005 under every null; system 3 is non-significant (*p* ∈ {0.13, 0.19, 0.23}). *α* = 0.01; minimum resolvable *p* = 0.00498 (200 replicates).

### Box 1 | What scJDO identifies

scJDO robustly identifies (empirically invariant across seeds and architectures):

- Relative operator structure (which genes co-vary in their sensitivity)
- Temporal ordering of operator regimes, e.g., sequential handoffs (recovered up to permutation on synthetic ground truth; matched Kendall τ = 0.97 on sigmoid handoffs, 22/24 seeds ≥ 0.9; 0.85 on sharp-step handoffs, 17/24 seeds ≥ 0.9, under operator-pattern matching; see Fig. 2b)
- Top-ranked genes associated with locally sensitive directions (though gene-level recovery is less precise than operator-level structure)

scJDO identifies the following only conditionally, and we treat them with corresponding caution:

- **Timing of local-sensitivity peaks**. Peak pseudotime is recoverable only when the underlying ordering is itself reliable; where pseudotime is poorly determined (e.g., near fate bifurcations, or where independent pseudotime estimators disagree), peak location is not robust and we do not make timing claims (see Discussion).
- **Concurrent (co-timed) archetype activations within a narrow pseudotime window**. The unregularized semi-NMF returns concurrent activations as spread peaks (matched concurrent-peak-pair prevalence 0.025 vs ground-truth 1.00). The Frobenius objective does not distinguish these solutions (residual 0.1101 for GT concurrent vs 0.1105 for spread-peaks, a ratio of 1.003). This is non-identifiability with respect to the objective: the choice is made by the prior, and the unregularized default’s implicit prior favors spread peaks. An opt-in `tv_lambda` temporal-smoothness penalty tips ALS toward the concurrent solution but with a bias–variance tradeoff: recovery of approximately half the concurrent signal (0.49 at *λ* = 10) coincides with destruction of sequential recovery on ground-truth-sequential systems at the same setting; the two operating points are incompatible on the current estimator. Concurrent-peak claims within a narrow window should be made only with the `tv_lambda` opt-in enabled, and cross-checked against sequential-recovery on a synthetic control.

scJDO is not designed to identify:

- Absolute biological Jacobians (exact entries of the regulatory matrix)
- Absolute eigenvalues (scale depends on pseudotime parameterization)
- Cluster-level rankings of which state is most unstable (prior-dependent; and see the Bifurcation-saddle limit below for the mechanism)
- Causal regulatory interaction strengths

Rather than recovering a unique underlying regulatory Jacobian, scJDO targets features of local dynamical structure that are reproducible across independently trained models and representations. Throughout this paper, “instability” refers strictly to the local mathematical sensitivity of the inferred transcriptomic drift field — the sign and magnitude of the dominant real eigenvalue of the local Jacobian — and not to physical instability or direct measurement of regulatory mechanism.

Stated affirmatively, the results below establish four things. Operator regime ordering is recoverable on synthetic ground truth up to the intrinsic permutation ambiguity of the decomposition, and is preserved under a mild temporal-smoothness prior on real data. Gene-level and directional operator structure shares significantly more information with an independent Jacobian method (Dynamo) than chance, despite that method resting on a different inferential prior — though gene-level recovery remains less precise than operator-level structure, and transcription-factor recovery is the weakest layer. Where the temporal axis is observed rather than inferred, operator handoffs are localisable in time and reproducible across decomposition settings. And saddle localisation is not recoverable from a drift field fit to snapshot density, with the post-saddle density-dominance signature visible across mathematically distinct families of readout.

We first establish what the estimator recovers on analytic ground truth and justify the latent-space choice (Fig. 2); then apply the framework to hematopoiesis (Fig. 3) and cross-validate against SpliceJAC and Dynamo (Fig. 4); characterize the bifurcation-saddle identifiability limit that constrains timing and cluster-level ranking claims for pseudotime-ordered data; and finish on a dense iPSC reprogramming time course with an observed experimental time axis that licenses the timing-resolved claim (Fig. 5), followed by a K562 Schrödinger-bridge feasibility demonstration (Fig. 6).

**Figure 3.**
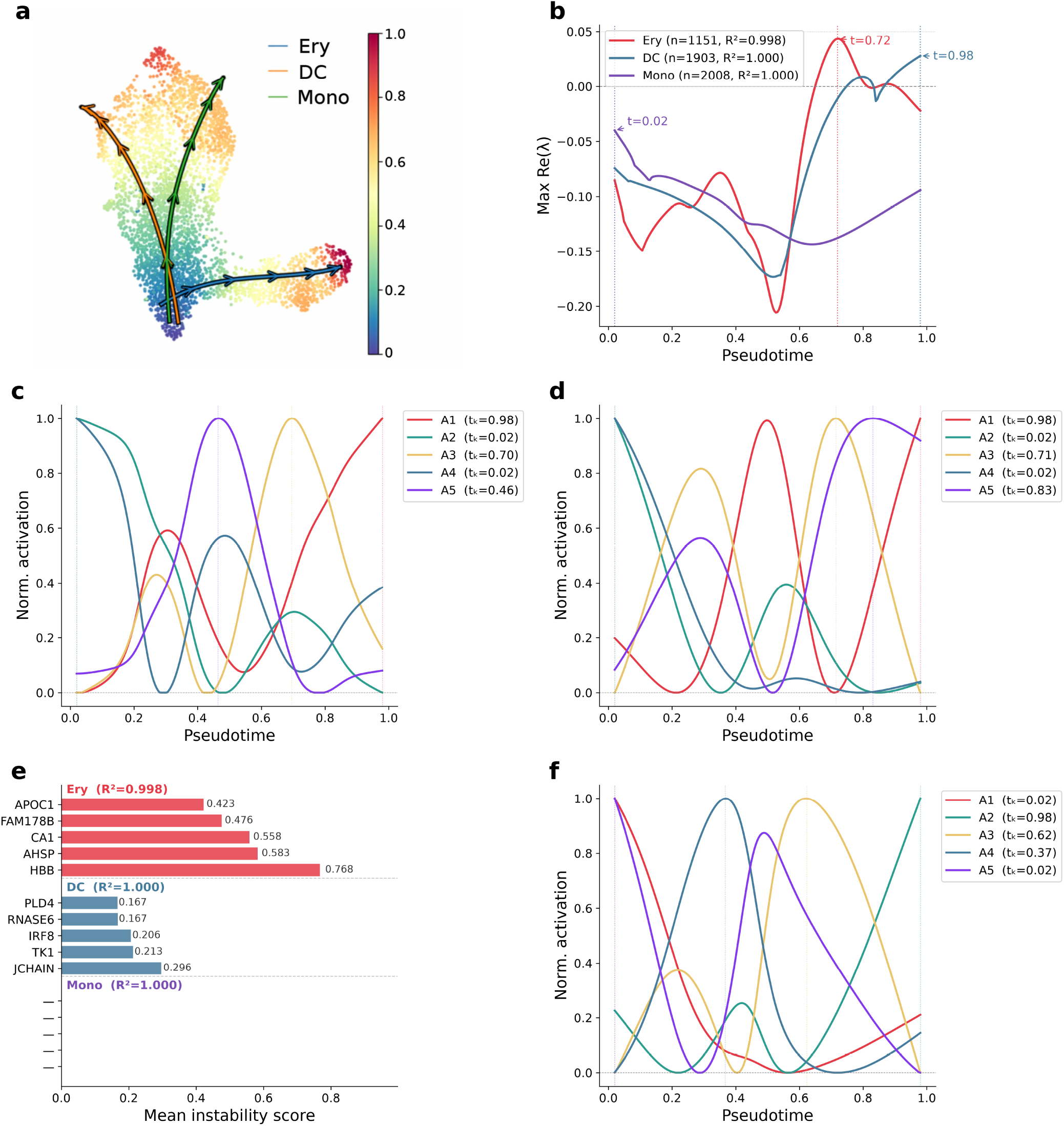
Branch-separated scJDO on the Palantir/Setty CD34+ bone marrow. 4,142 cells; 30-D Factor-Analysis; Palantir pseudotime; branch drift fields fit independently. **(a)** UMAP coloured by pseudotime with progenitor and terminal centroids. **(b)** Re(*λ*_max_) per branch: Ery (*n* = 1,151), DC (*n* = 1,903), Mono (*n* = 2,008). Mono stays below zero (informative negative). Peak-pseudotime values are annotated but not claimed as biological findings (Discussion §Bifurcation-saddle limit). **(c, d, f)** *K* = 5 archetype activation profiles for Ery, DC, Mono; component labels permute across branches (Methods §Cross-branch archetype matching). **(e)** Top instability genes: Ery — HBB, AHSP, CA1, FAM178B, APOC1 (GATA1 recovered without supervision as the top TF regulator); DC — TK1, JCHAIN, IRF8, CLSPN, C12ORF75; Mono — none recovered. *R*^2^ annotations on panels are the in-sample operator reconstruction fraction Ω (Methods), not drift-field predictive accuracy.

**Figure 4.**
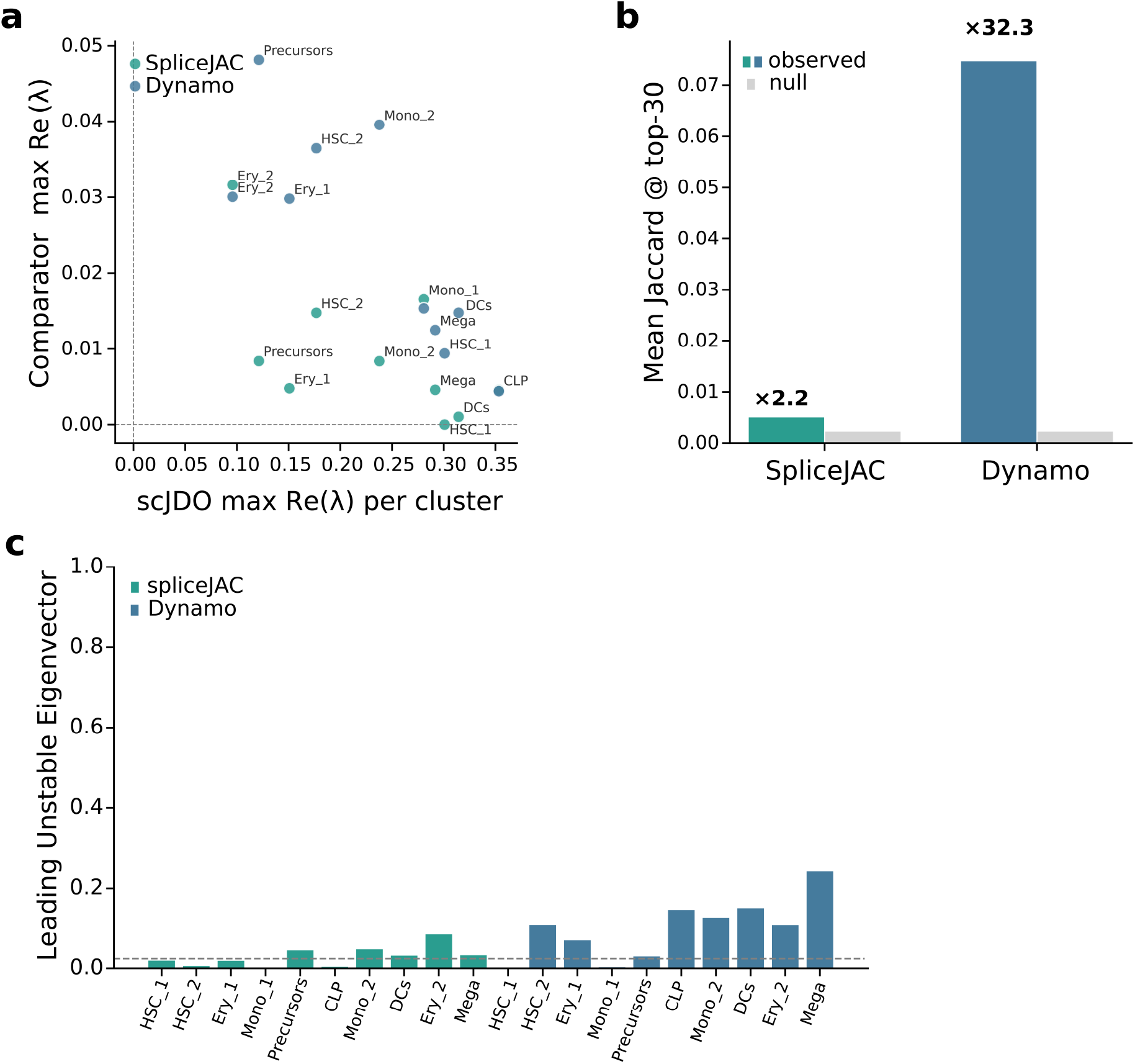
scJDO concordance vs SpliceJAC and Dynamo on bone-marrow CD34+. Ten shared clusters (HSC_1/2, Ery_1/2, Mono_1/2, Precursors, CLP, DCs, Mega); 6,482 shared genes. **(a)** Per-cluster Re(*λ*_max_) for scJDO vs each comparator; Spearman *ρ* = −0.77 (SpliceJAC) and *ρ* = −0.82 (Dynamo, *p* = 3.8 × 10^−3^) — systematic inversion consistent with the density-dominance mechanism (Discussion). **(b)** Mean Jaccard of top-30 transition genes: Dynamo 0.075 (32× random-gene null, ≈ 10× top-2000-HVG null); SpliceJAC 0.005 (2.2× random; 0.7× HVG). scJDO shares gene-level and directional structure with Dynamo while providing a distinct ordering. **(c)** Per-cluster absolute cosine of leading unstable eigenvector; null 95th ≈ 0.02. Dynamo above null in 8/10 clusters (mean. cos. = 0.100); SpliceJAC in 5/10 (0.030).

**Figure 5.**
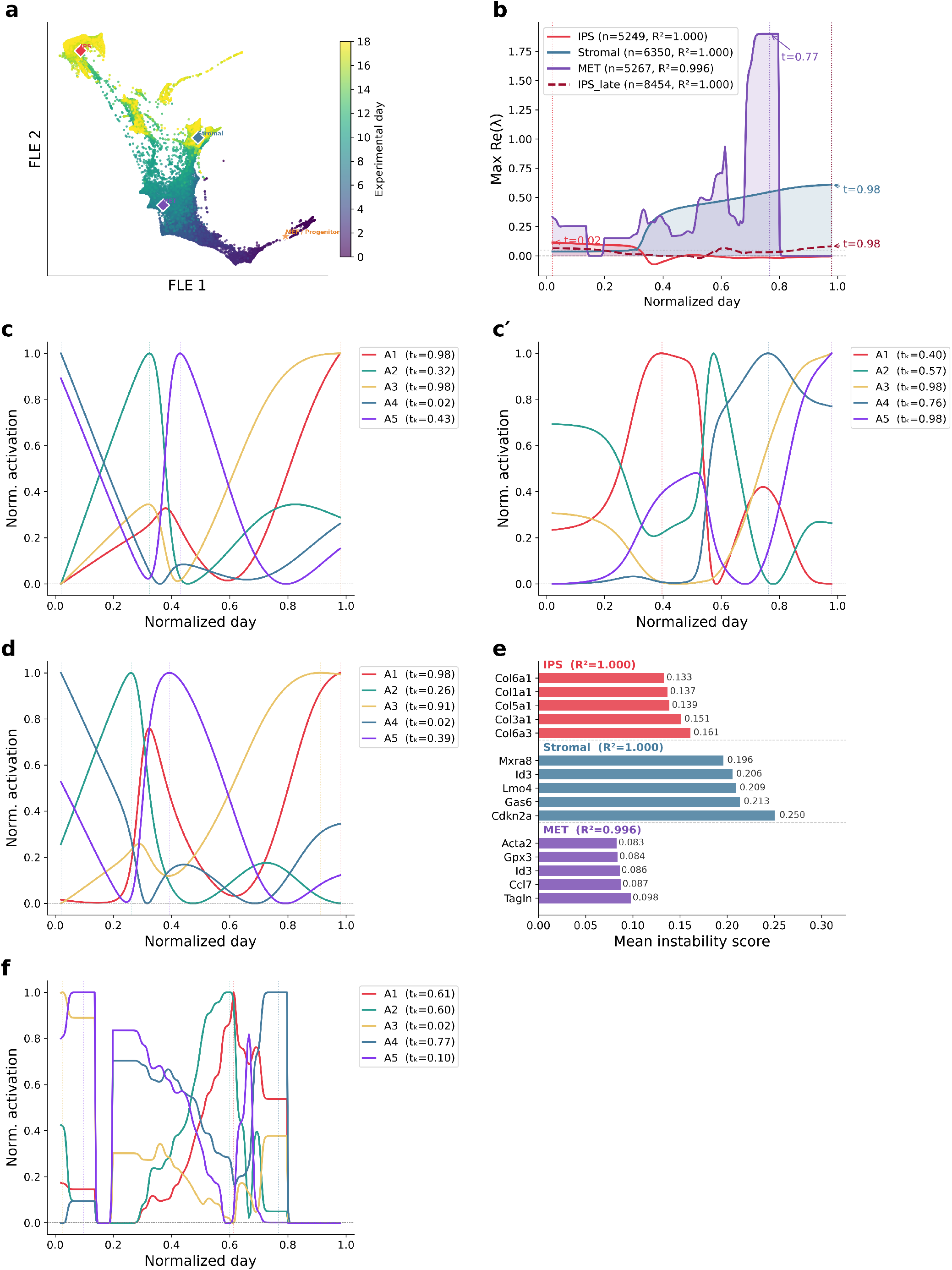
MEF-to-iPSC reprogramming (SCP295) — productive vs diverted operator handoffs. 24,999 cells across experimental days 0–18; 30-D FactorAnalysis; FLE embedding. Shown for both unregularised semi-NMF (Figure5_TV0.pdf) and the mild temporal-smoothness-prior control (Figure5_TV01.pdf, `tv_lambda = 0.1`); results are consistent between the two up to a permutation of component labels. **(a)** FLE coloured by experimental day, with MEF/progenitor and IPS/Stromal/MET centroids. **(b)** Re(*λ*_max_) per branch: IPS (*n* = 5,249), Stromal (*n* = 6,350), MET (*n* = 5,267), IPS_late (*n* = 8,454, day ≥ 4). MET exhibits a pronounced transient sensitivity peak. **(c, c’, d, f)** Per-branch archetype profiles (*K* = 5) for IPS, IPS_late, Stromal, MET. Productive branches (IPS, MET) execute a sequential handoff from an early **MEF-exit-like** archetype (*τ* ≈ 0.02, mesenchymal/ECM-exit genes) to a late pluripotency/metabolic-shift archetype (*τ* ≈ 0.6–0.75); Stromal maintains the early MEF-exit-like archetype while a distinct stress-associated program becomes active (Methods §Cross-branch archetype matching). **(e)** Top instability genes: IPS — Col6a3, Col3a1, Col5a1, Col1a1, Col6a1; Stromal — Cdkn2a, Gas6, Lmo4, Id3, Mxra8; MET — Tagln, Ccl7, Id3, Gpx3, Acta2 (top TF regulators include Trp53, Sp1, Nfkb1, Smad3, Rela). *R*^2^ annotations are Ω (see Fig. 3 note).

**Figure 6.**
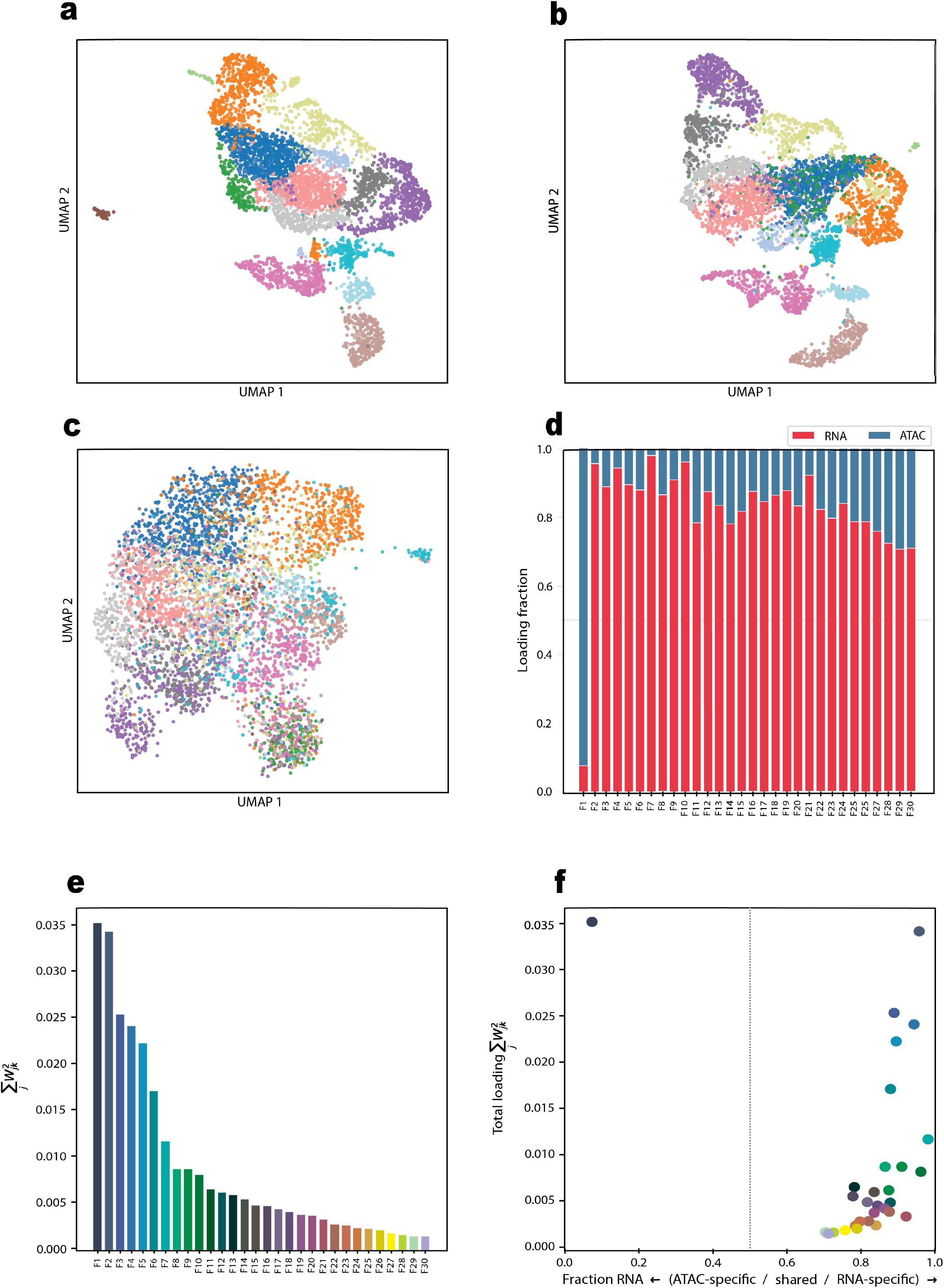
K562 CRISPRi Perturb-seq — per-perturbation Schrödinger bridges. 30-D FactorAnalysis; forward bridges from non-targeting-control (NT) to each of the three largest lncRNA-knockdown populations. **(a)** UMAP of NT (*n* = 1,238) with PVT1 (*n* = 135), MALAT1 (*n* = 126), and PSMA3-AS1 (*n* = 89) overlaid. **(b)** Forward-bridge Re(*λ*_max_) vs bridge time (NT → perturbed); peaks at *τ* = 0.97 with magnitudes +3.170 (PVT1), +0.644 (PSMA3-AS1), +0.311 (MALAT1). Magnitudes are not claimed as significant without per-target permutation (Fig. 6 validation). **(c, d, f)** Per-target *K* = 5 archetype activation profiles. **(e)** Top forward-instability genes: PVT1 — HNRNPA2B1, CSF3R, FAM178B, H1-2, TALAM1 (gene-level nomination only); MALAT1 — TOP2A, TPX2, CDCA8, UBE2S, CDC20; PSMA3-AS1 — TOP2A, H1-4, H4C3, TPX2, CDCA8. MALAT1 and PSMA3-AS1 converge on a shared mitotic/G2M signature consistent with the known cell-cycle-arrest phenotype of these knockdowns.

### Synthetic benchmarks validate the mathematical core and motivate Factor Analysis for operator inference

We first validated scJDO in synthetic systems with analytically specified dynamics, isolating method performance from biological complexity. In the bifurcating double-well benchmark, the learned drift field reproduced the true trajectory with low endpoint error (endpoint MSE = 0.0005) and assigned all simulated trajectories to the correct attractor, supporting the interpretation that scJDO learns a smooth continuous-time surrogate consistent with noisy observations. Along the trajectory, the estimated dominant Jacobian eigenvalue tracked the analytic eigenspectrum (eigenvalue correlation = 0.784) and recovered the stable-to-unstable sign transition, although exact gene-level projection of unstable directions remained approximate establishing early that scJDO is more reliable at the operator level than at single-gene resolution. The archetype-decomposition benchmark separated a temporal Jacobian sequence into three regimes corresponding to stable, transitional, and oscillatory dynamics (reconstruction error 0.228), supporting the interpretation that the decomposition step recovers distinct coordinated operator regimes when they are present. The Schrödinger-bridge benchmark [14] recovered endpoint-constrained transport close to the analytic target (*W*_2_ = 0.065).

A pipeline recovery probe on the same synthetic tensors quantifies how well the full pipeline (Jacobian → semi-NMF → archetype recovery) preserves the temporal-ordering signal. Because semi-NMF is ambiguous up to a column permutation of (*W*, *H*), we match recovered → ground-truth columns by cosine similarity on the operator matrices *H* (Hungarian assignment [16]; matching on *H* rather than the activation profiles *W* avoids the circularity of matching-on-activations then scoring activation ordering). After matching, mean Kendall τ across 24 seeds is 0.85 on sharp-step handoffs (17/24 seeds at τ ≥ 0.9) and 0.97 on sigmoid hand-offs (22/24 seeds at τ ≥ 0.9), with mean per-archetype cosine similarity on matched *H* pairs of 0.81 and 0.87 respectively — sequential-handoff structure is recoverable on synthetic ground truth up to the intrinsic permutation ambiguity. On concurrent overlapping-Gaussian activations the unregularized semi-NMF returns matched concurrent-peak-pair prevalence of 0.025 (vs ground truth 1.00). This is a basin-of-attraction property of the algorithm, not an identifiability limit of the data: initializing *W* at the ground-truth activations and running the same ALS converges to residual 0.1101 with matched concurrent-peak-pair prevalence 1.00 on all 8 seeds, whereas random-init cold starts land in a wider spread-peaks basin at residual 0.1105 (only 0.4% worse). The GT solution is a valid, slightly-better local minimum that random initialization does not reach; adding a total-variation temporal prior (`tv_lambda` opt-in, below) tips the algorithm toward it at a cost documented in the Failure Modes.

We then asked which latent representation best supports operator inference, benchmarking eight embeddings (PCA, Factor Analysis, ICA, TruncatedSVD, DiffMap [2], LDVAE [17], scVI [18], PLS) on bone marrow hematopoiesis using identical Palantir pseudotime and branch masks across all methods to isolate the effect of the representation. The choice is constrained before any benchmark is run. Because scJDO’s analytical object is a derivative, and because the implementation projects eigenvectors to gene space through a single fixed loading matrix, the latent map must be affine for that projection to be position-independent; under a nonlinear embedding the correspondence between a latent direction and a gene-space direction is the embedding’s own local Jacobian, which varies with position, so one global loading matrix no longer defines a consistent gene-space interpretation. This rules out scVI, LDVAE, DiffMap, and UMAP [19] for the fixed loading-based projection scJDO currently uses — not as a statement about nonlinear representations in principle, which could support local operator interpretation if their embedding Jacobians were explicitly propagated — and the benchmark bears this out: DiffMap returns no instability genes on any branch and is unscorable on specificity. Within the linear family, Factor Analysis (FA) models per-gene unique variances explicitly, which matches the heteroscedastic technical noise of scRNA-seq in a way that PCA’s isotropic residual assumption does not.

The peak-timing coordination metric that scJDO uses to score sequential-handoff structure was independently validated as a shape-agnostic positive control on four synthetic systems (sharp gated pulses, sigmoid handoffs, concurrent overlapping Gaussians, and unstructured curves; representative activation profiles in Fig. 2a). Across 24 independent seeds per system the metric approaches 1 on all three coordinated systems and remains at chance (≈ 0.5) on the unstructured system (Fig. 2b), and against three null models (circular shift, peak resampling, smooth randomisation) it reaches the minimum resolvable *p* < 0.005 on every coordinated system while remaining non-significant on the unstructured control (*p* ∈ {0.13, 0.19, 0.23}; Fig. 2c,d).

The benchmark is therefore a confirmation of this reasoning rather than its basis. On the synthetic operator-recovery benchmark (Supplementary Table 1) operator-recovery accuracy is comparable across PCA, ICA, and FA, with FA giving the tightest variance across noise levels; we do not read the accuracy comparison as discriminating among the three. On the Palantir/Setty [1] mouse-marrow sample FA returns high branch specificity — the Ery and DC top-20 instability gene sets have zero Jaccard overlap on this dataset, though this is a single branch pair on a single dataset at a fixed list length and should not be read as a general property. We also computed an aggregated composite score across marker enrichment, branch specificity, seed stability, timing-spread, and drift-field *R*^2^; we do not rest the choice of FA on this composite, because FA leads under only one of three defensible scoring configurations (PLS achieves the highest composite under the original specificity rule and again when the timing component is dropped; FA leads under the revised specificity handling). The full 8-method composite table under both specificity rules with and without the timing component is Supplementary Table 1, together with a discussion of scoring sensitivity; the qualitative picture (FA and PLS lead depending on rule; DiffMap is unscorable on specificity; the linear methods PCA / ICA / FA cluster together on synthetic operator recovery) is consistent across the table. PLS is supervised on branch labels and therefore functions as an upper bound rather than a default; scoring PLS on branch specificity is partially circular because PLS’s regression target contains branch information. We note that the “intermediate regularization” reading of FA’s suitability — enough denoising to stabilise drift-field derivatives, not so much smoothing that sharp transition-sensitive directions are attenuated — is a plausible mechanism consistent with our results but is not directly tested here; a monotone degradation of operator recovery from raw expression through linear to strongly nonlinear embeddings would be required to establish it. This choice was validated on hematopoiesis; generalisation beyond hematopoiesis is asserted rather than shown, and the reprogramming (Fig. 5) and multiome (Supp. Note) analyses use FA without a separately-validated out-of-domain embedding benchmark.

### Hematopoiesis reveals branch-specific local sensitivity

We applied scJDO to the Palantir/Setty [1] mouse bone-marrow sample (the 4,142-cell dataset from the Palantir tutorial, originally from Setty *et al*. 2019 CD34+ profiling), which spans differentiation from progenitors to erythroid, dendritic-cell (DC), and monocyte lineages. Using a 30-dimensional FA latent space and Palantir pseudotime (Fig. 3a), we inferred branch-separated drift fields and applied Jacobian analysis, which revealed branch-specific, non-uniform patterns of local sensitivity.

The erythroid branch (*n* = 1,151) gave the clearest result: a transient sensitivity regime (Fig. 3b) with top instability genes HBB, AHSP, CA1, and HBD — canonical erythroid maturation markers (Fig. 3e) — and GATA1 recovered without supervision as the top transcription-factor regulator; archetype activation profiles for this branch are shown in Fig. 3c. This is a particularly strong result given the documented field-wide bias across Jacobian methods toward recovering effector genes over transcriptional regulators due to TFs’ lower and noisier expression [20]. The biological interpretation of this branch rests on the recovered gene and regulator identity, which is robust across seeds, rather than on the precise peak pseudotime. Across nine tested configurations, the Palantir-vs-DPT branch-restricted Spearman ρ on the erythroid branch fell in the range 0.839–0.988, consistent with typical Palantir/DPT agreement on this dataset [1,2]. Because the two pseudotime estimators agree strongly on this branch, we decline peak-timing claims on the erythroid branch on the more general grounds discussed below rather than on pseudotime-estimator disagreement. Under our synthetic analysis, operator-timing recovery still depends on the interaction between the underlying ordering and the low-variance-transverse geometry near a branch point, so we report the erythroid result as a structural and gene-level finding and explicitly decline a peak-timing claim for it.

The DC branch (*n* = 1,903) exhibited a near-terminal sensitivity peak (Fig. 3b) with a mixed signature combining immune/DC-identity factors (JCHAIN, IRF8, IRF7, PLD4) and proliferative genes (TK1, RRM2) (Fig. 3e); archetype activation profiles for this branch are shown in Fig. 3d. Because this peak lies very close to the pseudotime boundary, we recommend confirming effective sample size (*n*_eff_) near the terminus and peak stability under bandwidth variation before treating its location as a primary claim; the proliferative component of this signature may additionally reflect the cell-cycle dominance discussed in the Supplementary Note, and should be assessed accordingly before biological interpretation.

In contrast, the monocyte branch (*n* = 2,008) did not exhibit a positive-eigenvalue sensitivity regime under the same criterion (Fig. 3b, flat below zero), and no instability genes or regulators were recovered (empty row in Fig. 3e; archetype activations in Fig. 3f). This negative result is informative: it suggests a comparatively buffered operator trajectory in the monocyte lineage in this dataset, and demonstrates that scJDO does not impose sensitivity peaks on all branches. Together, these branch-specific results illustrate how scJDO captures the molecular drivers of commitment where sensitivity is genuinely present, while remaining selective where the operator trajectory is buffered.

### scJDO shares operator-level structure with Dynamo but is distinct from splicing-derived Jacobians

To test whether scJDO’s learned operators reflect genuine biology rather than artifacts of the neural drift field, we compared it with the two most widely used single-cell Jacobian methods — SpliceJAC [8] and Dynamo [7] on the Setty 2019 [1] CD34+ bone marrow dataset, selected because it provides the spliced/unspliced layers these methods require. (A third method, scMomentum [21], was excluded from this comparison because it enforces a square Jacobian with identical regulators and effectors fitted cluster-wise via a Hopfield network, and requires source-code modifications to compute a Jacobian at all [20], making it unrunnable off-the-shelf for this comparison.) We compared three quantities: branch-level instability peaks, leading unstable-eigenvector directions, and top transition-gene lists.

Agreement with SpliceJAC was more modest than agreement with Dynamo across all three measures. On this dataset, the concordance pipeline operates on the 6,482 filtered genes shared by both methods and uses top-30 transition genes per cluster; the mean per-cluster Jaccard overlap is 0.0749 for Dynamo and 0.0051 for SpliceJAC (Fig. 4c). These are small absolute overlaps, and the informative quantity is the ratio to an appropriate null. We report the fold-over-null under two nulls: a random-gene null over the full 6,482-gene set and a more conservative null restricted to the shared top-2,000 HVG vocabulary that both methods actually draw from. Under the random-gene null, Dynamo overlap is 32× and SpliceJAC 2.2×; under the top-HVG null — the appropriate baseline given the shared HVG vocabulary — Dynamo’s mean Jaccard of 0.075 is approximately 10× the null and SpliceJAC’s 0.005 is 0.7× (i.e., SpliceJAC’s transition-gene overlap with scJDO is at or below the analytic null and not distinguishable from random draws over the shared HVG set). The full per-cluster breakdown — scJDO top-30, Dynamo top-30, their intersection, and null expectation for each cluster — is exported by the concordance notebook; a formatted supplementary table (Supp. Table 3) will accompany the journal submission. The leading unstable-eigenvector direction agreement (mean absolute cosine 0.100 for Dynamo vs 0.030 for SpliceJAC) was above the 95th-percentile null in 8 of 10 clusters (80%) for Dynamo vs 5 of 10 (50%) for SpliceJAC (Fig. 4b). We do not treat SpliceJAC as a reference standard for this comparison; we report the difference as informative about the two priors — splicing kinetics (SpliceJAC) versus cell-state geometry (scJDO) — rather than as a failure of either. Because Dynamo is built on a different inferential prior than scJDO, the shared gene-level and directional structure is evidence that this structure reflects the data rather than the assumptions of either method, not that either method is a ground-truth reference.

At the same time, scJDO and Dynamo disagreed on which branch carried peak instability (scJDO: CLP; Dynamo: Precursors), with the cluster-level instability ranking systematically inverted (*ρ* = −0.818, *p* = 3.8 × 10^−3^; Fig. 4a). We interpret this as a stronger statement than “prior-dependent”. The direction of the inversion is consistent with the account developed in the Bifurcation-saddle subsection below: if the dominant-eigenvalue readout tracks the density-dominant separation between already-committed fate populations rather than the low-variance transverse instability that defines a fate decision, then the cluster-level ranking is plausibly reading terminal-state variance rather than instability. While recent independent benchmarking validated Dynamo’s leading-eigenvalue transition detection against an analytical ground-truth Jacobian [20], that validation was performed on a driven, quasi-one-dimensional trajectory through saddle-node bifurcations — a regime with no low-variance transverse direction and no fate-decision saddle. That is precisely the committed-branch regime where our own Re(λ) readout succeeds. Dynamo is thus validated in the regime where the two methods agree, but the systematic inversion here suggests that at least one method’s cluster-level ranking is tracking a systematic artifact of its dominant-eigenvalue readout at fate decisions, which is precisely why we do not claim cluster-level instability rankings from either method. Because no ground-truth instability ranking exists for this dataset, and because scJDO’s gene- and regulator-level outputs on our biological datasets are consistent with established lineage biology, the concordance indicates that the shared structure reflects the data rather than the assumptions of either method, while scJDO provides a distinct, non-redundant operator-level view of hematopoietic sensitivity — with the ranking caveat named explicitly rather than left to the reader to reconstruct.

### Bifurcation-saddle limit: characterization and mechanism

The recurrent branch-point failure mode of density-based operator inference is important enough to warrant its own subsection with dedicated controls rather than a general concession in the Discussion. Recent independent benchmarking by Balubaid et al. [20] provides external corroboration for this limit: in their synthetic MET system, the analytical ground-truth Jacobian itself failed to detect the first of two transitions, establishing an upper bound on what any method could recover. That a true Jacobian fails to localize a bifurcation from this data class — and on a different bifurcation type (saddle-node) than our pitchfork — strongly suggests this is a general property of the modality rather than an artifact of one specific system.

We constructed a synthetic system specifically to test recovery of instability at a fate-decision saddle — the topology underlying our biological branching analyses. The system is a symmetric toggle-switch SDE with *α* ramped linearly through the pitchfork bifurcation (Methods). For the printed toggle system, the analytic pitchfork occurs at *α*_crit_ = 3^−1/4^(1 + 1/3) ≈ 1.013, corresponding to *τ*_crit_ ≈ 0.004. We therefore score saddle localization against an early window around the analytic crossing, *τ* ∈ [0, 0.054]. On this system, with analytically specified ground truth and oracle ordering, the dominant-eigenvalue readout does not recover the saddle location: the recovered peak in Re(λ) lands at *τ* ≈ 0.83, deep inside the committed-branch region, rather than near the analytic crossing.

We then tested whether alternative dynamical-systems readouts computed on the *same trained drift field* could recover the saddle where the dominant eigenvalue does not. Three principled alternatives target the transverse instability directly: the divergence tr(*J*) (which condenses/expands volume rather than the leading direction), the maximum real eigenvalue of the drift-tangent-projected Jacobian *P*_⊥_*JP*_⊥_ with 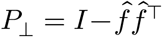 (which projects out the dominant flow direction), and the finite-time Lyapunov exponent (FTLE) [22] of the flow computed via `scjdo.atlas.flow_topology` (finite-difference flow-map Jacobian over an integration horizon of length 0.5 pseudotime units starting from each seed cell, then aggregated onto the pseudotime grid *τ* ∈ [0, 1] via the same Gaussian kernel used for the other readouts). The dominant-eigenvalue readout and the two readouts most directly aimed at transverse growth do not localize the analytic saddle: mean *τ*_peak_ = 0.83 for Re(λ), 0.85 for *P*_⊥_*JP*_⊥_, and 0.78 for FTLE. The divergence readout is sign-dependent, with peaks at *τ* ≈ 0.95 and *τ* ≈ 0.05 for the two signs; the early divergence peak lies at the upper boundary of the early scoring window. We therefore treat divergence as a boundary volume-change effect rather than a robust recovery of the transverse saddle instability. The failure of Re(λ), *P*_⊥_*JP*_⊥_, and FTLE supports the conclusion that the saddle limit is not a peculiarity of one scalar summary, while the divergence boundary effect cautions against overstating the negative as applying identically to every possible local readout.

To test whether the saddle limit is specific to eigenvalue-based readouts, and whether the density-dominance mechanism we invoke to explain the failure of the four eigenvalue-based readouts above has corroboration from a different direction on the same drift field, we additionally computed four operator-geometry descriptors via the windowed-Koopman backend [23] (Methods; `scjdo.tl.decompose_archetypes(method=`‘`koopman`’)): the Henrici departure-from-normality index [24]

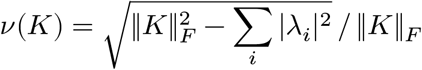

the reactivity ratio *σ*_max_(*K*)/*ρ*(*K*), the transient gain max *n* ≤ 30‖*K*^*n*^‖_2, and the eigenvector conditioning *κ*(*V*). Each is computed directly on the local reduced Koopman operator *K*_*τ* without a logarithm, so this family is immune to the log *λ* = log *λ* + 2*πik* branch ambiguity that constrains continuous-time rate estimates and is unaffected by any small *δτ* normalization artifact — a mathematically disjoint family from the four eigenvalue-based readouts above.

The four geometry descriptors do not behave uniformly. Henrici and reactivity — two measures of the same underlying departure from normality — both increase smoothly and monotonically across pseudotime, from *ν* ≈ 0.5 to *ν* ≈ 0.9 and from *σ*_max_/*ρ* ≈ 1.3 to ≈ 4.7, peaking at *τ* ≈ 0.90: the maximum-separation state of the two committed branches, not the analytic saddle at *τ*_crit_ ≈ 0.004 (Fig. S8b). An operator that is nearly normal at a saddle (transverse and along-flow directions comparably sensitive) and becomes progressively non-normal post-saddle as two committed branches diverge would produce a trend of this shape, so the Henrici and reactivity curves are consistent with the proposed density-dominance mechanism, computed via machinery that shares none of the eigenvalue-based readouts’ spectral construction. Confirmation across independent seeds would be required to move this from “consistent with” to a stronger inferential claim. Transient gain and eigenvector conditioning are noise-limited on this system, with argmax picking single-window numerical spikes from nearly-defective *K*_*τ*_ (interior peaks at *τ* ≈ 0.29 and *τ* ≈ 0.51; Fig. S8c); they do not cleanly test saddle localisation, but they do diagnose local eigen-fragility — a useful calibration for downstream users interpreting per-window Koopman eigenvectors. Summing across families: four eigenvalue-based readouts fail to localise the saddle (Fig. S8a), the log-free non-normality descriptors show a trend consistent with the density-dominance signature (Fig. S8b), and two descriptors are noise-limited (Fig. S8c). We describe these as mathematically distinct rather than statistically independent readouts: all eight derive from the same trained drift field, and the four eigenvalue-based readouts share much of their spectral construction.

A density-enrichment control was previously performed by simulating 5× the baseline cell density inside *τ* ∈ [0.117, 0.217] (same SDE, same drift-field training, two independent seeds, oracle ordering). Under the corrected analytic pitchfork location, this interval is a later post-saddle transition window rather than the analytic saddle window. We therefore treat this experiment as a later-window density-enrichment control, not as a definitive dense-sampling test at the analytic pitchfork. Under this enrichment, no readout shifts to the analytic early window: mean over 2 seeds *τ*_peak_ = 0.93 for Re(λ), 0.95/0.05 for tr(*J*), 0.86 for *P*_⊥_*JP*_⊥_, and 0.86 for FTLE (values reproduced in `scratchpad_run/saddle_dense_sampling.json`). Thus increasing local density in the later transition region does not rescue the saddle readout, but dense sampling centred exactly on *τ*_crit_ remains an explicitly untested follow-up.

These controls support the interpretation that the residual limit arises upstream of the scalar readout: a smooth drift field fit to snapshot density preferentially reproduces density-dominant separation between committed branches rather than the low-variance transverse instability that defines the saddle. The present evidence is strongest for the dominant-eigenvalue, transverse-projected, and FTLE readouts, and is further supported by the log-free geometry family: the non-normality descriptors show a trend consistent with the density-separation signature — monotonic growth peaking at the maximum-separation state — via mathematics that shares none of the eigenvalue-based readouts’ spectral construction. Divergence produces a boundary signal and should not be treated as definitive saddle recovery.

We emphasize that the density-dominance mechanism identified above is specific to the saddle regime and does not blanket-invalidate the operator-magnitude readout elsewhere. The regime where the artifact dominates is characterized by (i) an informative direction that is low-variance relative to the dominant density flow and (ii) a small, transient subpopulation of cells occupying that direction. Along a committed branch as in the erythroid transient sensitivity regime (Fig. 3) or the MET instability window (Fig. 5) — the sensitive direction is not low-variance relative to the local density flow, and the transiting cells are not a transient minority; the same Re(λ) readout therefore surfaces genuine gene-level biology at those locations even while it fails at the fate-decision saddle. A synthetic control that exhibits a non-bifurcating transient instability localized correctly by Re(λ) on a trained drift field would strengthen this scoping argument, and a matched null system with no transient instability would additionally test whether the bandwidth-selection objective can manufacture peaks. Our current synthetic controls (Systems 1a/1b) validate the decomposition step on ground-truth Jacobian tensors, but they do not pass through the drift-field training pipeline. We therefore flag this paired null-and-positive end-to-end control as a natural next test rather than a reported result. The remaining experimental levers are per-cell velocity information (metabolic labeling) that constrains the drift field directly, and lineage barcoding that measures the connectivity that pseudotime cannot recover at a saddle. This delineation directly informs how we frame the biological branching results below: we make claims at the level of operator structure and regime ordering, and we decline claims about the precise timing or ranking of branch-point instability.

### Dense reprogramming time-course reveals branch-specific operator handoffs

To assess whether scJDO’s operator regimes are robust to real time-course sampling rather than only snapshot-derived pseudotime, we applied the framework to a densely sampled iPSC reprogramming dataset (SCP295) [13], which captures the continuous transition from mouse embryonic fibroblasts (MEFs) to iPSCs over 18 days, with known branch points separating successful reprogramming from alternative fates such as stromal diversion. Critically, this dataset supplies an observed temporal axis — experimental day — rather than an inferred pseudotime, which is what enables the timing-resolved claims in this section that we decline to make for the pseudotime-ordered hematopoietic data.

The inferred drift field accurately captured the temporal progression of reprogramming when mapped onto the force-directed layout colored by experimental time (Fig. 5a). We analyzed four major branches: IPS (early successful trajectory), MET corridor, Stromal diversion, and IPS_late (late successful trajectory) (Fig. 5b). Because absolute eigenvalue magnitudes are not directly comparable across independently fitted branch drift fields, we report magnitudes only within the fitted branch model.

Within the MET corridor branch model, the dominant sensitive modes were dominated by mesenchymal and stress-associated genes (Tagln, Ccl7, Id3, Gpx3, Acta2; Fig. 5b, e, f), with Trp53, Sp1, Nfkb1, Smad3, and Rela among the top inferred regulators — consistent with a productive instability window during exit from the fibroblast/mesenchymal state. We note, however, that TF-level recovery is the weakest layer across all Jacobian methods [20], consistent with our posture that gene-level recovery is less precise than operator-level structure. We do not rank instability magnitudes across the four branches, because the drift fields are fit independently and the absolute-eigenvalue scales are not directly comparable across them. The precise MET-corridor peak pseudotime is bandwidth-dependent and should be interpreted as approximate: on a simplified reproduction of the MET-branch pipeline (3,344 MET-subset cells, 1,000 training epochs, no velocity-guidance prior; Supplementary Fig. S6) the supported peak τ (interior, *n*_eff_ ≥ 20 reporting filter for the sweep, distinct from the *n*_eff,min_ = 30 floor used in bandwidth auto-selection; Methods) shifts monotonically from 0.60 at *h* = 0.01 to 0.95 at *h* = 0.15, and only *h* = 0.01 returns a positive peak eigenvalue on that simplified pipeline. The full Figure 5 pipeline uses the auto-selected bandwidth on the complete branch and returns a positive-eigenvalue peak at τ ≈ 0.77; readers should treat the specific peak pseudotime as a bandwidth-conditioned estimate rather than an unambiguous data property, and any timing claim in downstream applications should be audited against a bandwidth-sweep sensitivity check on the exact same pipeline (Methods). The Stromal-diversion branch exhibited a distinct late instability regime marked by senescence-and stress-associated genes (Cdkn2a, Gas6, Lmo4, Mxra8; Fig. 5d, e) and enriched for NFκB/p53-associated regulators. Stromal diversion does not simply lack instability; it enters a qualitatively distinct late regime rather than completing the productive MET-to-iPSC operator transition. Early IPS cells showed a modest early sensitivity peak dominated by fibroblast/extracellular-matrix exit genes (Col6a3, Col3a1, Col5a1, Col1a1, Col6a1; Fig. 5c, e), while IPS_late cells showed a weaker terminal proliferative regime marked by cell-cycle genes (Birc5, Cenpf, Top2a, Ube2c, Cdca3; Fig. 5c’), reflecting expansion of successfully reprogrammed cells rather than a fate decision.

Archetype activation profiles further distinguished productive from diverted trajectories (Fig. 5c–f). Productive branches execute a sequential handoff from an early MEF-exit-like archetype (peak τ ≈ 0.02 on the MET corridor, top loadings on Col genes and mesenchymal-exit markers) through intermediate archetypes to a late pluripotency/metabolic-shift archetype (τ ≈ 0.61 on the MET corridor under TV=0; τ ≈ 0.69 on the IPS branch under TV=0.1). Because branch drift fields are fit independently and their semi-NMF component labels are arbitrary up to permutation and cone-ambiguity, we identify archetypes across branches below by their peak position and top-gene loadings rather than by component index (Methods §Cross-branch archetype matching); statements like “the same early MEF-exit archetype” therefore denote a functional identification, not a decomposition-level correspondence between one branch’s *A*_*k*_ and another’s. The productive-vs-stromal distinction is between whether this sequential handoff runs to completion (productive) or arrests at an early MEF-exit-like archetype (stromal diverted). The Stromal-diversion branch executed no such completion: it maintained an early MEF-exit-like archetype while a distinct stress-associated archetype became active. We deliberately avoid describing these as concurrent activations: Box 1 establishes that the unregularized semi-NMF used here is unreliable at resolving co-timed activations within a narrow pseudotime window, and we did not run the strong-TV concurrency diagnostic on this branch. The claim is the maintenance of the early archetype alongside a stress program, not their precise temporal coincidence. That the same estimator returns a sequential handoff for one branch and a maintained-archetype-plus-stress pattern for another is itself an internal control against a blanket separation bias of the unregularized semi-NMF.

To test whether the sequential early-to-late handoff was an artifact of the unregularized semi-NMF decomposition rather than a property of the reprogramming dynamics, we re-ran the entire Figure 5 pipeline with a mild total-variation smoothness prior on the semi-NMF activation profiles (`TV_LAMBDA = 0.1` opt-in in `jacobian_modes`, plumbed through `fit_drift_branches`; see Methods and `Figures_notebook/Figure4_tv01.ipynb`). The crossover occurs at approximately *t* = 0.69, while the late-archetype activation peaks later, at *t* = 0.72 without TV regularisation and *t* = 0.75 with mild TV regularisation. Thus TV=0.1 preserves the ordering and approximate timing of the handoff, although crossover time and peak time are distinct quantities.

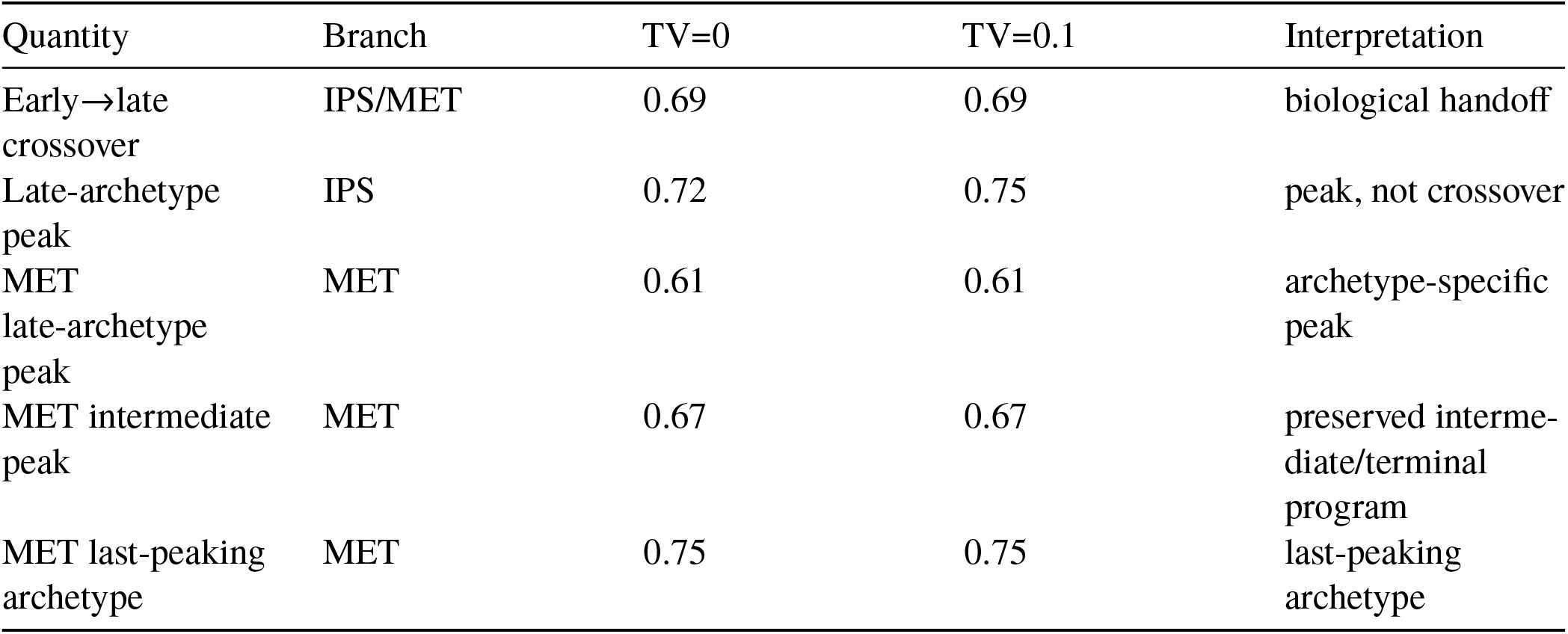

| Quantity | Branch | TV=0 | TV=0.1 | Interpretation |
| --- | --- | --- | --- | --- |
| Early→late crossover | IPS/MET | 0.69 | 0.69 | biological handoff |
| Late-archetype peak | IPS | 0.72 | 0.75 | peak, not crossover |
| MET late-archetype peak | MET | 0.61 | 0.61 | archetype-specific peak |
| MET intermediate peak | MET | 0.67 | 0.67 | preserved intermediate/terminal program |
| MET last-peaking archetype | MET | 0.75 | 0.75 | last-peaking archetype |

On the MET branch the archetype peak order is preserved between the two runs: peaks at 0.02 → (intermediate) → 0.61 → 0.67 → ≈0.75 in both. Semi-NMF’s permutation ambiguity re-indexes the archetypes (the same handoff appears under different component labels between runs, because the columns of *W* and rows of *H* can be permuted), but the temporal structure that the biological claim rests on is preserved (companion analysis: `Figures_notebook/Figure4_tv01.ipynb`). Independent synthetic controls confirm that TV=0.1 does not induce spurious sequentiality: on ground-truth-sequential Systems 1a and 1b of Fig. 2a, matched Kendall *τ* at *λ* = 0.1 is 1.00 (8/8 seeds) — TV=0.1 recovers sequential structure that is genuinely present without imposing it where it is not. The handoff at *t* ≈ 0.69 in the productive branches is therefore not a decomposition artifact of the unregularised default. Note the two operating regimes for `tv_lambda` are distinct: strong TV (e.g., *λ* = 10) is a diagnostic knob for concurrent-activation recovery — it tips ALS toward the concurrent minimum at the cost of destroying sequential recovery elsewhere; mild TV (*λ* ≈ 0.1) is a robustness check for sequential structure, preserving the sequential handoff both here and on the synthetic sequential systems. The two settings serve different diagnostic purposes and should not be read as “more vs less smoothing.”

This divergence in operator organization is present at earlier experimental time than the point of visible branch separation in the force-directed layout. Two caveats bound the interpretation. First, branch labels are derived from retrospective published annotations, so we frame this as a difference in operator organization between annotated branches rather than as a prospective, label-free early-warning signal. Second, because those annotations are themselves derived from transcriptional state, the comparison is not fully independent of the quantity being compared; a robustness check that perturbs the branch assignment near the annotation boundary, or that excludes the genes most predictive of branch label from the latent-space input, would be required to establish that the operator distinction is not partly inherited from the labeling. We flag this as a necessary follow-up rather than a completed control. Taken together, scJDO separates productive transition-state instability in the MET corridor from a distinct diverted stromal instability program, providing an operator-level account of reprogramming success versus diversion that complements differential expression and trajectory geometry.

### Per-perturbation Schrödinger bridges recover coherent regulatory programs in K562

While scJDO is primarily designed for developmental trajectories, its Schrödinger-bridge (SB) formulation [14] allows operator inference between defined source and target distributions, extending the framework to perturbational data where no natural pseudotime trajectory exists. We applied this to a K562 CRISPRi Perturb-seq dataset [15] in a 30-dimensional FA latent space, computing forward bridges from non-targeting control (NT) cells to the three lncRNA-knockdown populations with the largest cell counts: PVT1, MALAT1, and PSMA3-AS1 (Fig. 6a). This is presented as a feasibility demonstration in a regime where our central operator-magnitude readout is explicitly uninformative (see below); the value of the analysis is in the recovered gene-program content rather than in a magnitude ranking.

Each perturbation produced a distinct, biologically coherent program. The MALAT1 and PSMA3-AS1 bridges converged on a shared proliferation-arrest program (Fig. 6b, d, f) with strong, multiple-testing-robust enrichment for the G2M checkpoint (Hallmark enrichment [25]; FDR < 10^−12^, with the MALAT1 bridge reaching FDR ≈ 2 × 10^−17^ over 12 overlapping mitotic genes and the PSMA3-AS1 bridge reaching FDR ≈ 8 × 10^−13^ over 10 overlapping mitotic genes; Fig. 6 Validation, panel a), overlapping on canonical mitotic regulators including CDC20, CENPE, CENPF, and TOP2A (Fig. 6e). The PVT1 forward bridge nominated HNRNPA2B1 as its top forward instability gene (Fig. 6b, c, e); however, the associated pathway enrichment (PI3K–AKT–mTOR / mTORC1 axis) rested on a single overlapping gene and did not survive multiple-testing correction (FDR ≈ 0.64; Fig. 6 Validation, panel a). We therefore report the PVT1 program as a gene-level nomination only, and do not claim significant mTOR-axis enrichment for it; the only correction-robust enrichment in this analysis is the MALAT1 / PSMA3-AS1 G2M convergence.

Three points qualify the interpretation. First, while the recovered gene programs are coherent and distinct across perturbations, we assessed the significance of the bridge eigenvalue magnitude with a random-permutation null for the strongest signal only. For PVT1 — the strongest signal — the bridge eigenvalue was not distinguishable from a random null (empirical *p* = 0.143, 20 permutations; Fig. 6 Validation, panel b). We did not run the permutation control for MALAT1 or PSMA3-AS1; their significance rests on the pathway-level enrichment (G2M-Hallmark FDR ≈ 2 × 10^−17^ and ≈ 8 × 10^−13^ respectively), not on the eigenvalue magnitude. This is consistent with our caveat regarding static perturbation screens; the value of this analysis lies in the recovered program content, not in the instability magnitude. Second, because the gene programs are read from eigenvectors of the same operators whose magnitudes are indistinguishable from noise in the tested condition, an additional permutation null on program content (shuffling perturbation labels, rerunning the bridges, and checking whether the MALAT1 / PSMA3-AS1 G2M enrichment survives) is a natural next test to distinguish content-that-is-a-signal from content-that-is-a-consistent-shape-of-noise; we flag this as a caveat rather than reported evidence. Third, the mitotic enrichment recovered here reflects the genuine arrest phenotype of these perturbations — in contrast to developmental settings, where proliferation is a confound that must be regressed out at the level of latent-space construction (Supplementary Note). Note also that the MALAT1 / PSMA3-AS1 proliferation-arrest program is a known phenotype of those knockdowns and can be recovered by differential expression against non-targeting controls alone; the bridge adds a per-cell operator perspective on the transition, but the biological claim at the pathway level is not novel. Thus scJDO recovers interpretable, perturbation-specific operator programs even without a pseudotime ordering, with the explicit limitation that operator magnitudes are uninformative in this static regime.

## Discussion

scJDO treats the time-indexed sequence of local Jacobians as an explicit analytical object and decomposes it into recurrent operator archetypes with temporal activation profiles. Across synthetic systems, hematopoiesis, reprogramming, and perturbation screens, a consistent pattern emerges in what the method recovers reliably and what it does not, and that pattern is most usefully understood not as a list of separate limitations but as a single property of the underlying inference problem.

As established by Weinreb et al. [12], a drift field learned from a static transcriptomic snapshot is not uniquely determined by the data. A snapshot fixes the density of cell states but not the velocity field that generated it; many distinct drift fields are consistent with the same observed density, and any particular field is recovered only by imposing an additional assumption — splicing kinetics, a smoothness or gradient-flow prior, or a minimal-transport criterion. The Jacobian, and the local sensitivity we read from its spectrum, is therefore a derivative of an inferred quantity, conditioned on that prior. Recent independent benchmarking by Balubaid et al. [20] confirms this empirically, demonstrating that while inferred Jacobians recover behavioral properties, they struggle to recover structural ground truth. This has a direct and testable consequence: features of the operator that are determined by the data should be recovered robustly across different priors, whereas features determined by the prior should vary with it.

Our results separate cleanly along this line. The identity of the genes that dominate local sensitivity, the gene-space direction of the leading unstable mode, and the presence and ordering of operator regimes were robust — recovered consistently across linear embeddings, and sharing significantly more structure with an independent Jacobian method (Dynamo) than chance at the level of transition-gene overlap (mean Jaccard 0.075, approximately 32× a random-gene null over the 6,482 filtered genes shared by both methods, and ∼10× against the more conservative top-2,000 HVG-restricted null, which is the appropriate baseline given both methods draw from the same HVG vocabulary; SpliceJAC by contrast is at or below the top-HVG null) and eigenvector direction (above the null 95th-percentile in 8 of 10 clusters tested). That two methods built on different priors share significantly more gene-level operator structure than chance is evidence that this structure reflects the data rather than the assumptions of either method. By contrast, the quantities that depend on the inferential prior were not robust: the cluster-level ranking of which state is most unstable was systematically inverted relative to Dynamo (*ρ* = −0.818, *p* = 3.8 × 10^−3^), and exact instability magnitudes and peak pseudotimes were sensitive to choices upstream of the operator. We therefore frame scJDO’s biological claims at the level that survives — which genes and regulators drive local sensitivity, and the sequence of operator regimes — and explicitly decline claims about absolute magnitude, cluster-level instability ranking, or precise peak timing inferred from pseudotime alone.

This distinction matters for how our claims relate to work on the identifiability of full regulatory Jacobians. Recovering the complete operator — every entry of the gene-by-gene Jacobian — is a substantially harder estimation problem than recovering a low-dimensional set of reproducible operator descriptors. Negative results for full gene-by-gene Jacobian recovery therefore do not imply that lower-dimensional, reproducible operator summaries are unrecoverable. scJDO targets the descriptor level throughout, and we make no claim about full-matrix recovery.

The boundary is sharpest at fate decisions. The Bifurcation-saddle subsection in Results characterizes it directly: on the printed synthetic bifurcation, the analytic pitchfork occurs at *τ*_crit_ ≈ 0.004, and the dominant-eigenvalue readout peaks deep inside the committed-branch region rather than at the analytic crossing. Two readouts targeted to transverse growth — the transverse-projected spectrum *P*_⊥_*JP*_⊥_ and FTLE [22] — also peak deep inside the committed-branch region (*τ* ≈ 0.85 and *τ* ≈ 0.78). Divergence behaves differently: one sign peaks near the early boundary (*τ* ≈ 0.05) and the other near the terminal boundary (*τ* ≈ 0.95), making it a boundary volume-change signal rather than a robust recovery of the saddle. Four log-free operator-geometry descriptors from the windowed-Koopman backend give a complementary picture (Fig. S8): the non-normality measures increase monotonically along pseudotime and peak at the maximum-separation state (*τ* ≈ 0.90) rather than at the analytic saddle, a trend consistent with the proposed density-dominance mechanism, computed via mathematics disjoint from the eigenvalue-based readouts; transient gain and eigenvector conditioning are noise-limited and diagnose local eigen-fragility rather than testing saddle localisation cleanly. Because these descriptors avoid the log *λ* conversion entirely, neither the trend signal nor the noise signal can be attributed to Δ*τ* discretisation or the associated Nyquist branch. The failure is therefore not a quirk of Re(λ) or of any eigenvalue-based summary, and the geometry-family trend is consistent with the density-dominance account via mathematically disjoint machinery — though the failure signature is not identical across readouts. A later-window density-enrichment control (5× cell density in *τ* ∈ [0.117, 0.217], two seeds) does not shift any readout to the analytic early window, but under the corrected pitchfork location this control should not be described as dense sampling at the analytic saddle. We therefore interpret it as supportive evidence for density-dominance in the post-saddle separation regime, while treating dense sampling centered exactly on *τ*_crit_ as an untested follow-up. These controls support the interpretation that a smooth drift field fit to snapshot density preferentially reproduces density flow rather than the low-variance transverse instability at a saddle. Because the same density-dominance mechanism can operate at cluster level, we interpret the *ρ* = −0.818 inversion against Dynamo as a plausible instance of the dominant-eigenvalue readout reading terminal-state variance rather than instability, rather than merely as prior-dependence.

The bifurcation boundary is further clarified by the role of the time axis in our biological data. Where temporal order was observed rather than inferred — in the reprogramming time course, anchored to experimental day — scJDO localized an interpretable archetype handoff in time. Where temporal order was inferred from state geometry, recovery degraded at fate-decision saddles for the density-dominance reasons above rather than because independent pseudotime estimators disagreed: on the hematopoietic erythroid branch the two standard pseudotime estimators (Palantir [1] and DPT [2]) in fact agreed strongly (branch-restricted Spearman ρ in the range 0.839–0.988 across nine tested configurations), yet we still decline peak-timing claims for that branch because the density-dominance mechanism at branch points is upstream of the ordering. The contrast between the reprogramming and hematopoiesis settings is not incidental; it is the central evidence that the recoverable resolution of the method is set by the temporal information present in the data and by the local geometry near a decision boundary, not by the operator machinery: the biological committed-branch analyses produce interpretable transient sensitivity regimes when supplied with a well-resolved ordering, though end-to-end recovery of a known transient instability through the full learned-drift pipeline remains the untested control flagged above.

A related and underappreciated contributor to the difficulty at fate decisions is that the decision state is sparsely sampled to begin with. Because instability implies short residence time, cells pass through a saddle quickly and are correspondingly rare in any snapshot; this under-sampling is a property of the dynamics and is not remedied by sequencing more cells of the same kind. Our later-window density-enrichment control suggests that simply increasing density in the post-saddle separation region does not shift the learned readouts to the analytic saddle; dense sampling centered exactly on *τ*_crit_ remains an untested follow-up. It can be further compounded by standard preprocessing — quality-control filtering, doublet detection, and discrete clustering can each preferentially exclude or mislabel transcriptionally ambiguous transitional cells, which co-express competing programs and superficially resemble low-quality cells or doublets. We raise this as a caution for the field as much as for scJDO: where the transitional population is of interest, the cells that define it are precisely those most likely to be removed before analysis, and their recovery (e.g., via permissive branch-local QC and soft assignment) is one of the few interventions that can improve saddle sampling without a change of data modality.

These observations point to a concrete path beyond the present limits, and it is a path defined by the data modality rather than by a better estimator or a more principled scalar summary of the same drift field. Observed temporal information — dense experimental time courses, metabolic RNA labeling that supplies a per-cell clock, or lineage barcodes that measure rather than infer trajectory connectivity — directly constrains the velocity field that a snapshot leaves underdetermined. Our reprogramming result is preliminary evidence that observed time resolves the timing-dependent claims; a natural next test is whether single-cell metabolic labeling recovers operator timing where inferred pseudotime fails. Lineage tracing is, in principle, the strongest of these levers for the branch-point problem specifically, because it measures rather than infers the connectivity that pseudotime cannot determine at a saddle.

Read together, scJDO contributes an operator-level representation of single-cell dynamics — runnable on trajectory-resolved scRNA-seq datasets without splicing kinetics — and a characterization of the identifiability boundary of that representation. We regard these as a single contribution rather than a result and its caveats: the decomposition into temporal operator archetypes is, to our knowledge, a lens not previously available, and it is most useful when paired with an explicit account of the inference problem beneath it. Local dynamical sensitivity is recoverable at the level of structure and direction from snapshot data, while its magnitude and timing at fate decisions await the observed temporal information that current snapshot methods — scJDO among them — do not contain.

### Relation to existing dynamical approaches

scJDO occupies a distinct position in the single-cell analysis ecosystem. Transition-operator frameworks (e.g., CellRank [6], Palantir [1]) identify macrostates and fate probabilities, and continuous-time models (e.g., VeloVAE [26], neural ODEs [27]) learn trajectory geometry; these answer different questions from scJDO and are complementary to it (Table 1). At the level of operator decomposition, scJDO relates conceptually to Dynamic Mode Decomposition (DMD) [28] and Koopman operator theory [23], which identify recurrent linear regimes in dynamical systems. However, where global DMD fits a single linear operator to a time series, scJDO computes local nonlinear Jacobians per state and decomposes their temporal sequence. We additionally ship a *windowed* Koopman backend that fits local reduced Koopman operators on the vectorized Jacobian trajectory rather than on the raw expression time series (`scjdo.tl.decompose_archetypes(method=`‘`koopman`’)); this preserves the local-nonlinear-Jacobian core while exposing a spectral and geometry summary complementary to semi-NMF (see Methods and the toggle-switch geometry-readout controls above). The closest prior single-cell methods are the Jacobian frameworks SpliceJAC [8] and Dynamo [7], which compute local Jacobians and read off instability or regulatory structure per state. scJDO differs from these in three ways: it derives the Jacobian from cell-state geometry rather than splicing kinetics (removing the splicing-data requirement); it treats the time-ordered sequence of Jacobians as a primary object, decomposing it into recurrent archetypes rather than analyzing per-state snapshots; and it targets reproducible relative operator structure rather than causal regulatory entries. Our concordance analysis (Fig. 4) shows scJDO shares significant operator-level structure with Dynamo at the gene and direction level while providing a distinct ordering — establishing it as grounded but non-redundant.

**Table 1.**
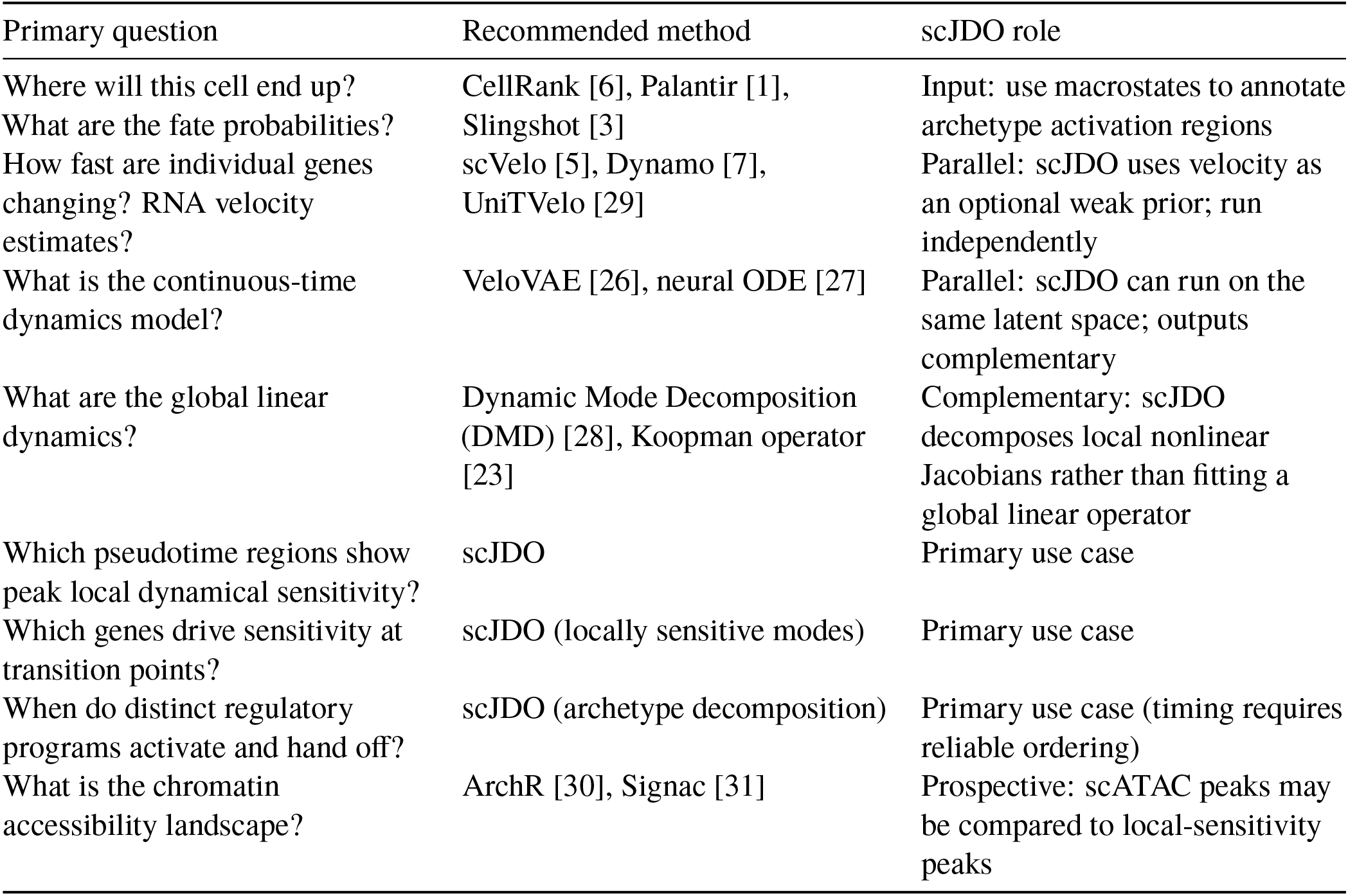
Decision guide for single-cell dynamics methods.

scJDO operates downstream of trajectory inference: it requires a pseudotime ordering as input and returns an operator-level summary of how local dynamical sensitivity evolves along that ordering. Users should first establish a biologically meaningful trajectory with their preferred method, then apply scJDO — bearing in mind that the reliability of any timing claim is bounded by the reliability of that ordering.

### Failure modes

Intellectual honesty about a method’s boundaries is as important as demonstrating its strengths. We identify conditions under which scJDO conclusions should be treated with particular caution or avoided.

#### Fate-decision saddles / bifurcation timing

As characterized in Results (“Bifurcation-saddle limit”), the dominant-eigenvalue readout and the two readouts most directly targeted to transverse growth (the transverse-projected spectrum and FTLE [22]) do not localize the analytic saddle in our synthetic bifurcation. Divergence produces a sign-dependent boundary signal rather than a robust recovery of the transverse saddle instability. Four operator-geometry descriptors from the windowed-Koopman backend (Henrici [24], reactivity, transient gain, eigenvector conditioning), which compute directly on the discrete-time operator and therefore avoid every log-based artifact, do not localize the pitchfork either — but the non-normality measures ramp monotonically to a peak at the maximum-separation state, a trend consistent with the density-dominance mechanism rather than a random miss (Fig. S8). A later-window density-enrichment control does not shift the readouts to the analytic early window, but under the corrected toggle-switch pitchfork location it should not be treated as dense sampling at the analytic pitchfork. Branch-point claims should therefore be made at the level of operator structure and regime ordering rather than precise timing or ranking of saddle instability, and the operator-magnitude readout at fate decisions is unlikely to be rescued by another local derivative of the same learned drift field without additional temporal or lineage information.

#### Static perturbation screens

scJDO is not appropriate for static perturbation screens lacking a biologically meaningful trajectory; in such settings pseudotime is arbitrary and the inferred operator magnitudes can become degenerate. This is directly observed in our K562 analysis, where gene programs were coherent but the operator magnitude for the strongest tested signal (PVT1) was indistinguishable from random (*p* = 0.143, 20 permutations; permutation control was run for PVT1 only, and the MALAT1 / PSMA3-AS1 significance rests on pathway-level enrichment rather than on eigenvalue magnitude).

#### Sparse or narrow pseudotime coverage; under-sampled transition states

scJDO requires sufficient cell density across the pseudotime axis to estimate a smooth drift field. Where cells concentrate at a few time points or large intervals are sparsely sampled, the drift field is poorly constrained and the resulting Jacobians reflect extrapolation rather than data. This is aggravated at fate decisions, where the transition state is intrinsically under-occupied and may additionally be depleted by standard QC, doublet removal, or clustering (Discussion). Near-terminal peaks (e.g., the DC branch) should be checked for effective sample size before interpretation.

#### Unreliable or method-dependent ordering

Because scJDO consumes a pseudotime ordering, timing claims are only as reliable as that ordering. Where independent pseudotime estimators disagree substantially, peak-timing claims should be withheld and interpretation restricted to gene/regulator identity and regime structure. (In our biological data the two standard estimators for the erythroid branch — Palantir [1] and DPT [2] — agreed strongly, branch-restricted Spearman ρ in the range 0.839–0.988 across nine tested configurations; we nevertheless decline peak-timing claims on that branch on the density-dominance grounds developed in the Bifurcation-saddle subsection.)

#### Externally derived branch or cluster labels

Where operator differences are compared across groups defined by pre-existing annotations, and those annotations are themselves derived from transcriptional state, the comparison is not fully independent of the quantity being compared. Our reprogramming analysis uses published branch annotations and is subject to this limitation; a label-perturbation or label-adjacent-gene-holdout control is required before an operator difference between annotated groups can be read as label-independent.

#### Concurrent-activation coordination within a narrow pseudotime window

The unregularized semi-NMF returns co-timed archetypes as spread peaks on synthetic ground truth (matched concurrent-peak-pair prevalence 0.025 vs 1.00). The Frobenius objective does not distinguish these solutions (residual 0.1101 for GT concurrent vs 0.1105 for spread-peaks, a ratio of 1.003). This is non-identifiability with respect to the objective: the choice is made by the prior, and the unregularized default’s implicit prior favors spread peaks. An opt-in `tv_lambda` penalty in `jacobian_modes` recovers approximately half the concurrent signal (0.49 at *λ* = 10) at the cost of destroying sequential recovery on ground-truth-sequential systems at the same *λ*; the two operating points are incompatible on the current estimator. Users making concurrent-peak claims within a narrow pseudotime window should enable the opt-in and cross-check against a synthetic sequential control at their chosen *λ*.

#### Cyclic or disconnected manifolds

The current implementation assumes a roughly tree-structured or linear progression. Cyclic trajectories (e.g., cell-cycle dynamics) and disconnected manifolds violate this assumption and may produce archetypes that reflect manifold topology rather than genuine dynamical regimes. Relatedly, in highly proliferative tissue the cell cycle can dominate the operator signal and must be addressed at the level of latent-space construction rather than gene-level regression alone (Supplementary Note).

#### Nonlinear latent representations

Because the Jacobian is a derivative in latent coordinates and its eigenvectors are interpreted only after projection through a single fixed loading matrix, that projection is position-independent only when the latent map is affine (Methods). Archetype identities are empirically robust to linear rotations, changes in latent dimension, and architectural variation. The current implementation should therefore not use nonlinear embeddings for gene-space operator projection unless their local decoder or embedding Jacobians are explicitly incorporated; this is a property of the fixed-loading projection as implemented, not a claim that nonlinear representations are inherently unsuitable for operator inference.

#### Systems dominated by cell–cell interactions

scJDO models cell-autonomous dynamics: the drift field depends only on the state of the individual cell and pseudotime, not on neighboring cells. Where fate is primarily governed by juxtacrine or paracrine signaling, microenvironmental gradients, or spatial coupling, the cell-autonomous model is misspecified and the inferred operators will not capture the dominant regulatory logic.

## Limitations

scJDO infers local linearized sensitivity of a learned drift field in latent space; it is not a direct measurement of physical regulatory interactions and should not be interpreted as a causal gene regulatory network. All biological language here is intentionally cautious: enrichment results are supportive rather than confirmatory, and operator structure is model-conditional rather than a ground-truth property of the underlying biology. Biological interpretation depends on projection to gene space and on external validation.

Because scJDO analyzes Jacobians of a learned drift field, its conclusions depend on how well that field is constrained by the available data and modeling assumptions. As developed in the Discussion, drift inference from snapshot data is fundamentally non-identifiable, so absolute magnitude, cluster-level ranking, and precise timing are prior-dependent and are not claimed; gene-level structure, operator direction, and regime ordering are the recoverable outputs. Coarse operator structure is more stable than fine-scale peak timing across reasonable aggregation bandwidths; peak location, and in some settings peak sign, can depend materially on bandwidth (see the MET bandwidth sweep in Results, where the supported peak shifts from τ ≈ 0.60 to τ ≈ 0.95 and only the narrowest bandwidth returns a positive peak eigenvalue on the simplified pipeline). Fine-scale archetype identity and peak timing should therefore be interpreted with the reported support diagnostics (Supplementary Fig. S2) and audited against a bandwidth sweep. All down-stream analyses use a 30-dimensional FA latent space; sensitivity of the biological results to this dimension is not reported here. The current implementation assumes cell-autonomous dynamics and does not model cell– cell communication. Inferred operators should be interpreted as transcriptomic dynamical surrogates rather than direct measurements of regulatory mechanism. Finally, while locally sensitive modes identify candidate perturbation-sensitive directions, causal conclusions will require targeted perturbations and prospective validation.

## Methods

### Latent representation selection

Because Jacobian inference computes local derivatives, the choice of latent representation is constrained by more than empirical performance. The Jacobian ∂*f*/∂*x* is defined in latent coordinates, and its eigenvectors are interpreted biologically only after projection to gene space. scJDO performs that projection with a single fixed loading matrix, and for a fixed loading-based projection the latent map must be affine: under a linear map with loadings *L*, a latent direction *v* corresponds to the gene-space direction *Lv* everywhere in the space. Under a nonlinear embedding the correspondence is the local Jacobian of the embedding itself, which varies with position, so one global loading matrix does not define a consistent gene-space interpretation and local warping confounds the differential operator. This is a constraint on the projection as implemented rather than a claim that nonlinear representations cannot in principle support local operator analysis — they can, if the embedding’s local Jacobian is explicitly propagated at each point, which the current implementation does not do. We therefore restrict scJDO’s default representation to linear embeddings, and treat nonlinear embeddings (UMAP [19], scVI [18], LDVAE [17], DiffMap [2]) as unsuitable for the fixed loading-based operator projection used here regardless of how well they preserve neighbourhoods or support visualization.

Within the linear family, Factor Analysis is the default on the grounds that its generative model assigns each gene an independent unique variance, which is the appropriate noise model for scRNA-seq — where technical variance is strongly gene-dependent — whereas PCA assumes isotropic residual variance and consequently allows high-noise genes to influence the leading components in proportion to their noise. This is a modeling argument, not a benchmark result, and we state it as the primary basis for the default.

We additionally benchmarked eight representations on hematopoiesis (Ery, DC, Mono lineages) as a confirmation. Operator-recovery accuracy on synthetic ground truth was comparable across the linear methods (PCA, ICA, FA), with FA showing the tightest variance across noise levels; DiffMap returned no instability genes on any branch. The composite score used to compare embeddings weights marker enrichment (30%), branch specificity (20%), seed stability (20%), timing spread (15%), and drift-field fit *R*^2^ (15%). Branch specificity is scored as 1 minus the mean pairwise Jaccard overlap of the top-N instability genes across branches; a branch that returns no instability genes (a correctly-buffered branch) contributes Jaccard 0 against every other branch (i.e., is treated as fully exclusive), and methods that return no instability genes for any branch are excluded from the composite ranking rather than scoring as maximally specific. Under this scoring, and with the timing component included, PLS achieves the highest composite (0.622); with the revised specificity handling FA moves to first (0.729); dropping the timing component (aligning the composite with our declared posture on timing claims) returns PLS to first (0.732) with FA second (0.706). We report all three configurations because the composite does not consistently favour FA, and we do not use it to justify the default. The full 8-method table under both specificity rules with and without timing is Supplementary Table 1.

The framework accepts any continuous latent space, and users are free to substitute another linear representation. We do not claim FA is universally superior for single-cell analysis; we claim that the affine-map requirement is structural for Jacobian projection, and that among linear representations FA’s noise model is the better match for scRNA-seq. This was validated on hematopoiesis; generalization to other tissues is asserted rather than validated out-of-domain, and users applying scJDO to substantially different tissues should re-run the embedding benchmark for their own data.

### Data preprocessing and latent-space embedding

Raw gene-expression matrices were normalized to counts per 10,000, log-transformed, and restricted to the top 2,000 highly variable genes (Scanpy v1.9.3 [32]; Seurat flavor). Dimensionality reduction used Factor Analysis with 30 components for all datasets, selected by the benchmarking described above. The embedding benchmark evaluated representations across dimensions 10, 30, and 50, with 30 components providing the best balance of denoising and local differential-structure preservation; the operator-recovery comparison among embeddings was performed at matched dimensionality. Sensitivity of the downstream biological results to the choice of 30 components was not systematically assessed and is noted as a limitation.

### Temporal coordinate inference and trajectory sampling

Main biological analyses used the trajectory method most appropriate for each dataset: Palantir [1] (v1.0.1) for the Palantir/Setty [1] mouse bone-marrow sample; diffusion pseudotime (DPT) [2] on a k-nearest-neighbor graph (*k* = 15; Euclidean distance in FA space) for robustness comparisons; experimental day as the ordering variable for SCP295 reprogramming. Slingshot [3] (v2.4.0) was used as an additional sensitivity check. DPT and Palantir can produce differently distributed pseudotime orderings, and we quantify their cross-method agreement per branch (rank correlation) as a diagnostic of ordering reliability; the adaptive kernel-windowing procedure (below) is sensitive to pseudotime density. For temporal alignment across methods and runs, activation profiles were aligned using dynamic time warping (DTW).

### Relation to velocity-based fate mapping

Velocity- and transition-operator methods [4–7] infer macrostates, terminal basins, and fate probabilities by constructing transition operators informed by RNA velocity. scJDO instead targets local operator structure by learning a smooth drift field and computing time-indexed Jacobians along inferred progression. Unless otherwise stated, scJDO results are reported without using CellRank-derived macrostates for archetype definition; macrostate annotations may be overlaid for visualization only. Velocity guidance, when used, is treated as a weak directional prior.

### Drift-field parameterization and learning

Cellular dynamics are modeled as *dx*(*t*) = *f* (*x*(*τ*), *τ*) *dt* + *σ*_drift_ *dW*_*t*_, where *τ* ∈ [0, 1] is the **biological pseudotime** (observed experimental day for SCP295; Palantir pseudotime for the Palantir/Setty [1] mouse bone-marrow sample; DPT for robustness checks) and *σ*_drift_ is the SDE diffusion scale.

#### Distinctions maintained throughout

We keep four objects separate: (i) biological pseudotime *τ* (the input ordering, computed by an external method); (ii) the **denoising noise scale** *σ* used in the diffusion-score-matching training objective (a nuisance scale over which we average, unrelated to *τ*); (iii) the **learned score field** 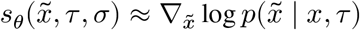— the network’s output during training, targeting the conditional perturbation score at pseudotime *τ*; and (iv) the **surrogate drift field** *f*_*θ*_(*x, τ*) whose Jacobian is the analytical object of the paper. Objects (iii) and (iv) are related but not identical: *f*_*θ*_ is constructed at inference from a hybrid architecture combining a *σ* → 0 evaluation of the score component with a residual deterministic head (see “Surrogate drift field” below), so the Jacobian analyzed downstream is *not* the raw denoising direction from the score net.

#### Score-matching objective (score component of *f*_*θ*_)

The score component of the drift network is trained via conditional denoising score matching [10,11]. For each training triple (*x, τ*, *σ*) with *σ* drawn from a log-uniform distribution over a fixed range, we form a perturbed sample 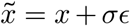 with *ϵ* ∼ *N*(0, *I*_*d*_) and minimize

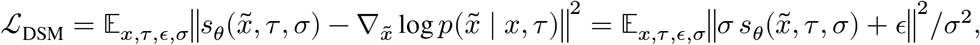

where the second equality is the standard equivalence to noise prediction [10]. This targets the **conditional perturbation score** 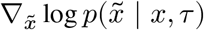, not the marginal data-density score 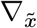 log 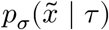. Under mild regularity the two agree in expectation over *x* at each 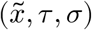; we use the conditional form for training stability and marginalize over *σ* at inference.

#### Surrogate drift field for operator analysis

The drift field whose Jacobian is the object of interest is

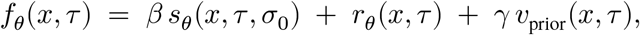

combining the score component (evaluated at a small nuisance scale *σ*_0_ → 0^+^; scaled by *β*), a **residual deterministic head** *r*_*θ*_ trained jointly with a pseudotime-gradient consistency term that pulls the field along the observed / inferred temporal ordering, and an optional **velocity-guidance prior** *v*_prior_ from RNA velocity [4,5] (weight *γ* ≥ 0; *γ* = 0 disables it). The residual head is what converts a denoising direction (which by itself tends to point back toward the data manifold) into an ordering-consistent developmental drift; the pseudotime-gradient term is a soft consistency penalty rather than a hard constraint. Crucially, while the score component *s*_*θ*_ is the gradient of a scalar potential (and thus its Jacobian is a symmetric Hessian), the residual head *r*_*θ*_ is unconstrained and breaks this symmetry, allowing the full surrogate drift *f*_*θ*_ to capture non-equilibrium rotational or oscillatory dynamics. When we speak of “the drift field” for Jacobian analysis, we mean *f*_*θ*_ evaluated at the small nuisance scale *σ*_0_, not the raw score component alone. We do not claim that *f*_*θ*_ uniquely recovers the biological drift; scJDO uses *f*_*θ*_ as a smooth, prior-defined **surrogate drift** whose local derivatives summarize reproducible geometry-conditioned sensitivity along the supplied ordering, and the identifiability boundary of that surrogate (Bifurcation-saddle Results subsection, Box 1, Failure Modes) is inherited by every downstream operator claim.

#### Architecture and training

The drift *f*_*θ*_ was parameterized by a time-conditioned multilayer perceptron (four layers, width 256, SiLU activations) with residual skip connections, FiLM-style time conditioning [33] of both the score and residual heads on (*τ*, *σ*), and spectral normalization [34] on the final layer. The drift-field composition *f*_*θ*_(*x, τ*) = *β s*_*θ*_(*x, τ*, *σ*_0_)+*r*_*θ*_(*x, τ*)+*γ v*_prior_(*x, τ*) uses *β* = 0.1 (score-component scale) and *γ* = 2.0 (velocity-guidance strength) as the defaults for every analysis reported here; the pseudotime-gradient velocity prior enters as this structural component of the surrogate drift rather than as a separate loss term. The training objective is ℒ_DSM + *α*_control ℒ_control with *α*_control_ = 10^−3^, where 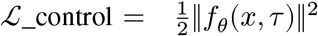 is a small control-energy penalty preventing runaway drift magnitudes. These three numerical weights (*β* = 0.1, *γ* = 2.0, *α*_control_ = 10^−3^) fully determine the surrogate drift whose Jacobian is the paper’s analytical object; the seed and training-set specification are additionally recorded per analysis in `analysis/configs/` and in the header of each per-figure notebook. Sensitivity to *γ* was checked by rerunning the MET branch with *γ* = 0 (velocity-guidance disabled) in the bandwidth-sweep supplement, which preserves the qualitative peak structure. Optimization used AdamW [35] (learning rate 2 × 10^−4^, weight decay 10^−4^), batch size 1,024, gradient clipping at 1.0, and cosine annealing, for 5,000 epochs on an NVIDIA A100 GPU. The nuisance noise scale *σ* is log-uniform on [10^−3^, 5 × 10^−1^] during training and *σ*_0_ = 10^−3^ at inference.

#### Operator reconstruction fraction (Ω)

The scalar exposed as `adata.uns[key][`“`r2`”] by `scjdo.tl.fit_drift` is the in-sample *operator reconstruction fraction*, 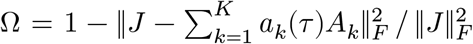, evaluated at the same pseudotime grid on which the temporal Jacobian tensor *J* was aggregated. It quantifies how well the *K* archetype basis reconstructs *J* on the training grid and nothing else. In particular, it is *not* a drift-field fit quality, *not* a lineage-progression score, and *not* a held-out generalisation score, so we do not report it in the biological Results and warn users against interpreting it as such. Held-out drift-field validation was not performed and is flagged as follow-up work.

### Jacobian inference and eigenanalysis

Local Jacobians were computed via automatic differentiation in PyTorch (v2.1.0): *J*(*x, τ*) = ∇_*x*_*f*(*x, τ*). We performed eigen-decomposition using `torch.linalg.eig` without symmetrization. Modes were classified by the maximum real eigenvalue: we classify modes as stable (real eigenvalue ≤ −0.04), plastic (in between), or sensitive (real eigenvalue ≥ +0.04), with thresholds selected to match the observed spectral gap between clearly-stable and clearly-sensitive regimes on the training dataset. These absolute thresholds are parameterization-dependent and are used only for descriptive classification within a fitted branch model, not for cross-branch magnitude comparison. Eigenvectors were projected to gene space using FA loadings.

### Temporal Jacobian tensor construction and archetype decomposition

Because local Jacobian estimates depend on the temporal support used for averaging, scJDO estimates a continuous operator trajectory using kernel-weighted pseudotime neighborhoods rather than treating fixed bins as independent observations. For each pseudotime grid point *τ*, per-cell Jacobians are averaged with Gaussian weights:

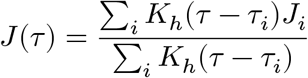

where *K*_*h*_ is a Gaussian kernel with bandwidth *h* and *τ*_*i*_ is the pseudotime of cell *i*. The bandwidth is selected automatically by maximizing a composite score that balances bootstrap reproducibility of the dominant-eigenvalue profile, instability-peak contrast, peak localization, and a minimum effective sample size (*n*_eff,min_ = 30). This floor is a constraint on bandwidth selection and is distinct from the *n*_eff_ ≥ 20 interior filter used when reporting supported peaks in the MET bandwidth sweep (Results). Because peak contrast is part of the selection objective, the reported peaks are not independent of the smoother choice; we treat the monocyte negative (a branch where the same objective returns no peak) as one control against the concern that the objective produces peaks by construction, and recommend a bandwidth-sweep sensitivity check (report peak pseudotime and contrast as a function of *h*) for any timing claim that a downstream user makes with a different dataset. A designed control — a matched pair of synthetic systems, one with no transient instability and one with a known localized non-bifurcating instability, both passed through the complete pipeline including automatic bandwidth selection — would test this more directly than the monocyte negative and is flagged as a next step. The effective sample size 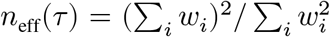 is retained as a diagnostic of local operator support and reported alongside instability profiles. This widens the temporal neighborhood in sparsely sampled regions, preventing visually sharp but under-supported peaks. The legacy fixed-window scheme (100 overlapping windows, 80% overlap) is retained as an optional sensitivity mode. Reported peak pseudotime values reflect the kernel-weighted estimate; timing resolution is limited by local pseudotime density and approximate values are reported accordingly.

Smoothed Jacobians were stacked into *J* ∈ ℝ^*T*×*d*×*d*^, unfolded into 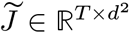, and projected into a lower-dimensional operator space. The primary decomposition used semi-nonnegative matrix factorization (semi-NMF [9]; *M* ≈ *W H, W* ≥ 0, *H* signed), with singular value decomposition (SVD) as a fallback when semi-NMF did not converge. *K* = 5 was fixed across all seven branches of both datasets (Ery, DC, Mono; IPS, Stromal, MET, IPS_late) to **standardise decomposition rank and model capacity** across branches, because per-branch cross-validated reconstruction curves are flat over *K* ≈ 4–6 on every branch. Fixing *K* standardises rank; it does **not** induce component correspondence across independently decomposed branches. Because semi-NMF is ambiguous up to column permutation (and, more broadly, up to a *Q*-cone ambiguity), archetype indices from independently fitted branch drift fields are not comparable — MET’s *A*_1_ is not the same object as Stromal’s *A*_1_. Cross-branch archetype comparisons in Results §Reprogramming are therefore made by peak position and gene content rather than by component index (see “Cross-branch archetype matching,” next). Optimisation alternates a closed-form least-squares update for the signed factor *H* and a column-wise non-negative least-squares update for *W* (Lawson–Hanson [36] via `scipy.optimize.nnls`, giving monotone descent of the Frobenius objective), with five random restarts; the best restart by final reconstruction error is retained. Post-convergence, the factors are rescaled so each row of *H* has unit *ℓ*_2_ norm, resolving the diagonal scale ambiguity of the (*W, H*) factorisation. An optional `tv_lambda` argument enables a total-variation penalty on the temporal derivatives of *W* (proximal-gradient inner loop with soft-thresholded first differences); this is not used in the main analyses (which use `tv_lambda = 0`, i.e., no temporal prior) and is provided in the package as an opt-in for analyses where concurrent-peak claims within a narrow pseudotime window are the biology of interest. Synthetic controls (Fig. 2b, and §Synthetic benchmarks Results) show that sequential-recovery quality is preserved at `tv_lambda ≈0.1` and destroyed at `tv_lambda ≥ 1`; users enabling the opt-in should audit their setting against these controls.

#### Cross-branch archetype matching

Branch drift fields are fit independently and decomposed independently (Ery/DC/Mono for hematopoiesis; IPS/Stromal/MET/IPS_late for reprogramming), so semi-NMF component indices are not comparable across branches (see K-selection paragraph above). Cross-branch statements in Results (“the same MEF-exit archetype”, “an early MEF-exit-like archetype”, etc.) are identifications by *peak position and gene content*, not by component index. Specifically, we call two archetypes cross-branch-matched when (i) their peak *τ* values fall within a single semi-window of one another (the same coarse pseudotime bin used for reporting peaks in Results) and (ii) their top-30 gene loadings overlap sufficiently to license a common functional label (e.g. mesenchymal/ECM-exit for “MEF-exit”). No programmatic matching (Hungarian [16] on operator cosine, mutual-nearest-neighbours on *H*-columns, etc.) is applied — the identifications are made by inspection of Fig. 5b–f and the per-branch instability-gene CSVs. A programmatic cross-branch matching pipeline (matching by cosine on *H*-columns, Hungarian-assigned, with a shuffled-branch null) is flagged as a follow-up. Readers should therefore treat cross-branch archetype identities as functional labels, not component-level correspondence.

### Bifurcation-saddle test system and alternative readouts

The bifurcation system (v3 of the synthetic benchmark) is a symmetric toggle-switch SDE *dx* = [*α*(*t*)/(1 + *y*^4^) − *x*]*dt* + *σ dW*, *dy* = [*α*(*t*)/(1 + *x*^4^) − *y*]*dt* + *σ dW*, with *α* ramped linearly from *α*_min_ = 1 (single stable symmetric fixed point) through *α*_crit_ = 3^−1/4^(1 + 1/3) ≈ 1.013 (pitchfork bifurcation) to *α*_max_ = 4. The analytic saddle crossing is at *τ*_crit_ = (*α*_crit_ −*α*_min_)/(*α*_max_ −*α*_min_) ≈ 0.004. Cells are generated by Euler– Maruyama integration with *σ*_SDE_ = 0.20; a random duration per cell distributes *τ* approximately uniformly over [0, 1]. Toggle-space coordinates are projected to 200-D observation space via a random linear projection with *σ* = 0.3 observation noise, then reduced to 20-D by PCA.

Four alternative readouts of the trained drift field are computed on the same kernel-aggregated pseudotime grid used for the baseline Re(λ): (1) divergence tr(*J*) (both signs, since a saddle may present as either sign depending on which direction dominates); (2) the maximum real eigenvalue of *P*_⊥_*JP*_⊥_ where 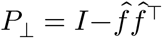 is the projector orthogonal to the drift tangent (targeting instability transverse to the dominant flow); (3) the finite-time Lyapunov exponent (FTLE) [22] computed via `scjdo.atlas.flow_topology.compute_ftle` (finite-difference flow-map Jacobian; integration horizon of length 0.5 pseudotime units starting from each seed cell; 50 Euler steps over that horizon; finite-difference step *δ* = 10^−3^). The per-cell FTLE values are then Gaussian-kernel aggregated onto the same *τ* ∈ [0, 1] grid used for the other readouts, so reported FTLE peak locations refer to positions on that grid rather than to the integration horizon. Peak localisation is scored against an early window around *τ*_crit_, *τ* ∈ [0, 0.054]. The baseline saddle-readout tabulation was computed from a single seed (`SEED=42`); the density-enrichment control (described next) uses two independent seeds. The density-enrichment control oversamples inside the [0.117, 0.217] interval by a factor of 5 (adding cells whose pseudotime is drawn uniformly from that interval on top of the baseline uniform coverage). Under the corrected pitchfork location, this interval is a later post-saddle transition window rather than the analytic saddle window; we therefore treat it as a later-window density-enrichment control rather than as dense sampling at the analytic pitchfork. Two seeds are a small sample and this negative should be treated as suggestive rather than definitive at this seed count.

Four additional readouts derived from the windowed-Koopman backend [23] are computed on the same trained drift field and the same pseudotime grid: (5) the Henrici departure-from-normality index [24]

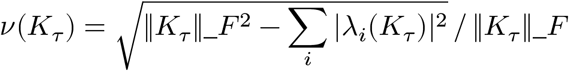

bounded in [0, 1] and zero iff *K*_*τ*_ is normal; (6) the reactivity ratio *σ*_max_(*K*_*τ*_)/*ρ*(*K*_*τ*_), bounded below by 1 and tight iff *K*_*τ*_ is normal; (7) the transient gain 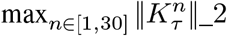, capturing short-horizon amplification that asymptotic eigenvalues can miss for non-normal *K*_*τ*_; (8) the eigenvector conditioning 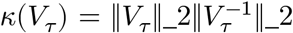 of the local eigenvector matrix. Descriptors (5) and (6) both measure departure from normality and are not independent of one another; we report both because they are conventional in different literatures, but treat their agreement as one line of evidence rather than two. Each descriptor is computed on the local reduced Koopman operator *K*_*τ*_ ∈ ℝ^*r*×*r*^ fit by extended DMD [23] with identity observables on the sliding-window {(*z*(*τ*_*i*_), *z*(*τ*_*i*+1_)) : *τ*_*i*_ ∈ *W*_*τ*_} pairs of the vectorized Jacobian trajectory *z*(*τ*) = *P*_*r*_vec(*J*(*τ*)) (window half-width 8 grid steps, Tikhonov ridge 10^−4^, `whiten=False` metric; the reduced-space metric echo is stored in the diagnostics dict so numbers and metric always travel together). Because these four descriptors are computed directly on the discrete-time operator without a logarithm, they are immune to the log *λ* = log *λ* + 2*πik* branch ambiguity that constrains log *λ*/Δ*τ* continuous-time rate estimates, and they are unaffected by any small Δ*τ* normalization artifact that a log /Δ*τ* conversion introduces. Peak localisation uses the same interior-argmax and ±0.05 tolerance as the eigenvalue-based readouts; the derivation and equations are given in `KOOPMAN.md` §7, and the exact script (`analysis/supp_saddle_localization/saddle_koopman_geometry.py`) imports the toggle-switch construction from the eigenvalue-based `saddle_readouts.py` so the ground-truth SDE, seed, latent dimension, drift training regime, and pseudotime grid are identical across the eight readouts.

As with the eigenvalue-based tabulation, the geometry descriptors were computed on a single seed (`SEED=42`). The monotonic Henrici and reactivity trends are smooth across the full pseudotime grid and unlikely to reflect seed noise, but confirmation across independent seeds — reporting distributions of *τ*_peak_, *ν*(*K*), and *σ*_max_(*K*)/*ρ*(*K*) rather than single-realisation curves — remains an untested follow-up, and is the control that would determine whether the transient-gain and eigenvector-conditioning spikes reflect a general noise limitation or an artifact of this realisation. On this system, the four descriptors do not behave uniformly: Henrici and reactivity vary smoothly and monotonically along pseudotime with interior peaks at the maximum-separation state (*τ* ≈ 0.90), so their argmax is a genuine structural feature consistent with the density-dominance signature rather than a random miss; transient gain and eigenvector conditioning show sharp single-window spikes from nearly-defective *K*_*τ*_ (interior argmax at *τ* ≈ 0.29 and *τ* ≈ 0.51), and are more usefully read as diagnostics of local eigen-fragility than as scalar saddle-localisation summaries. We report all four and separate their interpretation in Fig. S8.

### Schrödinger-bridge formulation

For the K562 CRISPRi Perturb-seq dataset [15], we employed a Schrödinger-bridge (SB) formulation [14] to regularize the drift field. The SB objective minimizes the Kullback–Leibler divergence to a reference Wiener process, subject to matching the empirical marginal distributions at the initial (non-targeting control) and final (perturbed) time points. This was implemented with an alternating Sinkhorn and score-matching algorithm, anchoring the inferred trajectory to the observed biological endpoints. Forward bridges were computed independently for each perturbation (PVT1, MALAT1, PSMA3-AS1) using NT cells as the source. Statistical significance of bridge eigenvalue magnitudes was assessed against a random null by permutation (reported as empirical *p*); the permutation control was executed for PVT1 only, and the significance of the MALAT1 and PSMA3-AS1 results rests on pathway-level enrichment (see Results). Pathway enrichment of bridge instability-gene sets used Hallmark gene sets [25] with Benjamini–Hochberg FDR correction; we report FDR-corrected values throughout and treat enrichments with FDR > 0.05 as non-significant.

### SCP295 iPSC reprogramming analysis

We analyzed *n* = 24,999 cells drawn as a stratified subsample of the SCP295 upload (Schiebinger et al. 2019 [13]) from serum and Dox reprogramming conditions; the stratified subsample is produced by `Figures_notebook/build_scp295_h5ad.py`. Four branch assignments — IPS, MET corridor, Stromal diversion, and IPS_late — were derived from published cell-set annotations; because those annotations are themselves derived from transcriptional state, operator comparisons across them are not fully independent of the labeling, and a label-perturbation control is flagged in Results as a required follow-up. The primary root was defined at day 0. Drift fields were fit independently for each branch in the FA latent space. Experimental day, normalized to [0, 1], was used as the temporal ordering variable.

### Concordance analysis (Setty CD34+)

For comparison with SpliceJAC [8] and Dynamo [7], we used the Setty 2019 [1] CD34+ bone marrow dataset, which provides spliced/unspliced layers required by both comparator methods. SpliceJAC and Dynamo Jacobians were computed per their published pipelines. scJDO operators were aligned to comparator clusters, and concordance was assessed by (i) branch-level maximum real eigenvalue and its cross-method rank correlation, (ii) absolute cosine similarity of leading unstable eigenvectors per cluster against a null threshold, and (iii) mean Jaccard overlap of top-30 transition genes against a random-gene null. The concordance pipeline itself operates on the 6,482 filtered genes shared by both methods, and the conservative null restricts the random draw to the shared top-2,000 HVG universe that overlaps the working gene set; the top-HVG restriction is a null construction rather than a pipeline gate. We note that a random-gene null is a permissive baseline for methods drawing from the same top-HVG axis on the same dataset; the top-HVG null produces a smaller fold-enrichment but does not change the qualitative agreement pattern. Absolute overlaps are small (mean per-cluster Jaccard 0.075 for Dynamo), so we report raw overlap alongside fold-over-null throughout; the full per-cluster breakdown — each method’s top-30 list, their intersection, and null expectation — will accompany the journal submission as Supplementary Table 3.

### Supplementary Note — cell-cycle handling in proliferative tissue

In highly proliferative lineages, the cell cycle is the highest-amplitude coordinated transcriptional program and can dominate the inferred operator. We found that regressing cell-cycle (S/G2M) scores from gene expression after latent-space construction is insufficient: the proliferation signal persists because it is encoded in the manifold geometry of the learned drift field. On the 10x Genomics E18 mouse-brain multiome dataset, the uncorrected joint FA latent does carry the cell-cycle signal directly — factor 1 has | Pearson *r*| = 0.50 with the G2M score and factor 2 has |*r*| = 0.32. Despite this manifold-level CC dominance, in the ExcNeuron branch of this dataset the top-9 instability genes are neurogenic markers (uncorrected: Cacna2d3, Col19a1, Neto1, Grik1, Npas1, Reln, Galntl6, Thrb, Kctd8; CC-regressed: Raly, Gnai2, Ndufa11, …) rather than the mitotic set (Cdc20, Ccnb1, Cenpf, …) sometimes reported in proliferative-lineage instability analyses. We therefore recommend the pre-integration correction as a defensive default and describe the correction as a robustness measure rather than a rescue: the invariance of canonical developmental marker ordering under the correction (Supplementary Table) supports both variants of the E18 pipeline as biologically valid. Concretely, we regress cell-cycle scores from both RNA and ATAC modalities before integration and latent-space construction; on the E18 mouse-brain multiome dataset, applying this correction preserves the neurogenic top-9 ExcNeuron ordering and leaves canonical developmental marker ordering invariant. We therefore recommend cell-cycle correction at the level of latent-space construction as a standard defensive preprocessing step when scJDO is applied to proliferative tissue, and we report this as a methodological caveat rather than a validation claim.

## Author Contributions

**David Redd**: Conceptualization, Methodology, Software, Formal analysis, Investigation, Writing — original draft, Writing — review & editing.

**Sam Green**: Methodology, Software, Validation, Data curation, Visualization, Writing — review & editing.

**Tommy W. Terooatea**: Conceptualization, Supervision, Project administration, Funding acquisition, Writing — review & editing.

## Data Availability

The mouse bone-marrow sample dataset used for Figure 3 is the 4,142-cell dataset from the Palantir tutorial (from Setty *et al*. 2019 [1]); it is available at https://github.com/dpeerlab/Palantir under `data/marrow_sample_scseq_counts.h5ad`. The Setty 2019 [1] CD34+ bone marrow dataset (used for the Figure 4 concordance analysis) is available from its original publication. The SCP295 iPSC reprogramming dataset [13] is available via the Single Cell Portal. The K562 CRISPRi Perturb-seq dataset used here is the genome-scale lncRNA sublibrary from Replogle et al. 2022 [15]; we analyzed non-targeting control cells plus the PVT1, MALAT1, and PSMA3-AS1 knockdown populations. The 10x Genomics E18 Mouse Brain Fresh 5k Multiome dataset (used in the Supplementary Note) is available at 10xgenomics.com. All accession numbers and data sources are listed in Supplementary Table 2, and `Figures_notebook/DATA.md` documents every external data source used by the notebooks.

## Code Availability

The scJDO package is publicly available at https://github.com/manarai/scJDO underan open-source license, and includes the drift-field estimation, Jacobian inference, archetype decomposition, Koopman geometry, and Schrödinger-bridge functionality described here. Analysis notebooks and configuration files used to generate the figures in this manuscript are available from the corresponding author upon reasonable request.

